# Small Molecule Screening in Zebrafish Embryos Identifies Signaling Pathways Regulating Early Thyroid Development

**DOI:** 10.1101/551861

**Authors:** Benoit Haerlingen, Dr. Robert Opitz, Isabelle Vandernoot, Dr. Achim Trubiroha, Pierre Gillotay, Nicoletta Giusti, Dr. Sabine Costagliola

## Abstract

**Background:** Defects in embryonic development of the thyroid gland are a major cause for congenital hypothyroidism in human newborns but the underlying molecular mechanisms are still poorly understood. Organ development relies on a tightly regulated interplay between extrinsic signaling cues and cell intrinsic factors. At present, however, there is limited knowledge about the specific extrinsic signaling cues that regulate foregut endoderm patterning, thyroid cell specification and subsequent morphogenetic processes in thyroid development.

**Methods:** To begin to address this problem in a systematic way, we used zebrafish embryos to perform a series of *in vivo* phenotype-driven chemical genetic screens to identify signaling cues regulating early thyroid development. For this purpose, we treated zebrafish embryos during different developmental periods with a panel of small molecule compounds known to manipulate the activity of major signaling pathways and scored phenotypic deviations in thyroid, endoderm and cardiovascular development using whole mount *in situ* hybridization and transgenic fluorescent reporter models.

**Results:** Systematic assessment of drugged embryos recovered a range of thyroid phenotypes including expansion, reduction or lack of the early thyroid anlage, defective thyroid budding as well as hypoplastic, enlarged or overtly disorganized presentation of the thyroid primordium after budding. Our pharmacological screening identified BMP and FGF signaling as key factors for thyroid specification and early thyroid organogenesis, highlight the importance of low Wnt activities during early development for thyroid specification and implicate drug-induced cardiac and vascular anomalies as likely indirect mechanisms causing various forms of thyroid dysgenesis.

**Conclusions:** By integrating the outcome of our screening efforts with previously available information from other model organisms including *Xenopus*, chicken and mouse, we conclude that signaling cues regulating thyroid development appear broadly conserved across vertebrates. We therefore expect that observations made in zebrafish can inform mammalian models of thyroid organogenesis to further our understanding of the molecular mechanisms of congenital thyroid diseases.

## Introduction

Thyroid hormones (TH) are critically important regulators of vertebrate growth and development (1). The thyroid gland is the only known site of *de novo* synthesis of TH in vertebrates. The minimal functional units of thyroid tissue are thyroid follicles consisting of a single layer epithelium of thyroid follicular cells (TFC) enclosing a luminal cavity that is filled with a proteinous matrix containing TH precursors (2, 3). Although thyroid tissue differs in shape, size and anatomical position between vertebrate species, these variations in thyroid anatomy appear to have no major impact on thyroid functionality (3). The key requirement for normal thyroid function is the formation of sufficient amounts of functional thyroid follicles to meet physiological demands for TH synthesis.

The thyroid is the anterior-most organ derived from the embryonic gut tube in vertebrates and its morphogenesis is a complex developmental process comprising a sequence of conserved morphogenetic events (2–4). The initial event in TFC development is the specification of thyroid precursor cells (TPC) within a restricted domain of the anterior endoderm. Molecularly, TPC are characterized by co-expression of a unique combination of transcription factors including orthologs or paralogs of human *NKX2-1, PAX8, FOXE1* and *HHEX* (5–7). This TPC population constitutes the thyroid anlage, formed as early as embryonic day 8.5 (E8.5) in mouse and around E20 in human (8, 9). TPC represent a committed cell state but functional differentiation genes are not yet expressed and TPC are not capable of TH synthesis. Current concepts hold also that the size of this population of committed cells is likely one key determinant of later organ size (6).

Morphologically, the earliest evidence of thyroid organogenesis is the formation of the thyroid placode recognizable as a thickening of the ventral foregut epithelium, commonly positioned at a level between the first and second pharyngeal arch in a region where the cardiac outflow tract (OFT) is in close contact with the pharyngeal endoderm (8, 10, 11). The placodal thickening rapidly transforms into a true organ bud that eventually detaches from the pharyngeal floor followed by a relocation of the thyroid diverticulum towards its final adult position. The thyroid primordium then expands by proliferation and attains a species-specific shape. Along with the initiation of folliculogenesis, thyroid functional differentiation genes are induced, encoding among others for thyroglobulin, thyroid-stimulating hormone receptor, thyroid peroxidase and sodium-iodide symporter (2). This induction also marks the transition of TPC into terminally differentiated TFC capable of TH synthesis. Thyroid organogenesis is complete in mice at E16.5 (8) and around the 11^th^ week of gestation in human (12).

Although the morphological events of thyroid organogenesis have been outlined in much detail in diverse model organisms (8, 13–16), the understanding of the specific molecular mechanisms regulating cell and tissue dynamics during these processes remains at best preliminary (3, 6, 16). Organ development relies on the tightly regulated interplay between extrinsic signaling cues and cell intrinsic factors (i.e. lineage-specific transcription factors). While core aspects of the transcription factor network regulating thyroid cell development have been experimentally dissected in mouse and zebrafish (7, 17–20), there is still a paucity of data on the extrinsic signaling cues that regulate foregut endoderm patterning, thyroid cell specification and subsequent morphogenetic processes (21). From recent reviews on these topics (6, 21, 22), it becomes apparent that current knowledge is largely based on a patchwork collection of experimental observations made in diverse experimental systems and species. This makes it challenging to formulate robust hypotheses on the role of distinct signals for specific morphogenetic processes and to infer the potential contribution of perturbed signaling to the etiopathology of developmental thyroid diseases such as congenital hypothyroidism due to dysgenesis (2, 6, 21).

Accordingly, there is a need for systematic analyses to further our understanding of the specific role of common signaling pathways during thyroid organogenesis. Suitable test systems to tackle this problem should meet some important demands. First, a recurrent theme in the developmental biology of endoderm-derived organs is that adjacent mesodermal tissues play a critical role as a source of inductive and permissive signals for organ specification and differentiation (23–25). With regard to thyroid development, fate mapping and tissue recombination experiments in various species revealed that pre-cardiac anterior lateral plate mesoderm and the OFT-forming cardiac mesoderm are likely major sources of signaling cues instructing early thyroid development (10, 26). Such complex tissue-tissue interactions are difficult to replicate in classical *in vitro* systems arguing for the need of whole organisms as experimental systems. In addition, current models of endoderm patterning and organ specification highlight that mesoderm-endoderm interactions can be transient and that specific signaling cues can have diverse and even opposite regulatory activities depending on the temporal context (27–29). Accordingly, experimental models preferably should offer the ability to have temporal control over the phenotype. In this respect, chemical genetic approaches based on small molecules targeting specific biological pathways can provide distinct advantages over classical genetic models.

Zebrafish embryos represent one such powerful *in vivo* model system to investigate cellular and molecular mechanisms of thyroid organogenesis combining the tractability of worm or fly models with physiological and anatomical characteristics of higher vertebrates (30). Morphogenetic events during early thyroid development are well conserved from fish to men (7, 11, 13–15, 18, 22, 31) and the zebrafish model offers several salient features that are particularly attractive for larger-scale chemical genetic studies on organ development (32). In contrast to the intrauterine development of mammalian embryos, zebrafish embryos develop externally which greatly facilitates experimental manipulations and permits monitoring of developmental processes and pathological deviations in real time. Addition and removal of bioactive compounds at specific developmental stages offers temporal control of embryonic signaling activities in order to characterize phenotypes within specific developmental contexts. The high fecundity of zebrafish permits generation of thousands of developmentally synchronized embryos that can then be housed conveniently in well plates or small petri dishes. Combined with small size, optical transparency and rapid development of embryos, these advantageous properties permit the realization of *in vivo* screening approaches requiring large numbers of test organisms (32, 33).

In this study, we harnessed advantages of the zebrafish model to perform a series of pharmacological screens investigating the role of major signaling pathways during early thyroid organogenesis. For this purpose, we treated embryos during defined developmental periods with a panel of small molecules known to modulate specific signaling pathways and performed comprehensive phenotypic analyses with respect to thyroid morphogenesis and cardiovascular development. A systematic assessment of drugged embryos recovered a range of different thyroid phenotypes including expansion, reduction or lack of the early thyroid anlage and the thyroid primordium, defective thyroid budding as well as overtly disorganized anatomical presentation of the thyroid primordium. Our results highlight key roles for fibroblast growth factor (FGF) and bone morphogenetic protein (BMP) signaling during thyroid specification, demonstrate the importance of low Wnt/β-catenin signaling activities for early morphogenetic events and implicate drug-induced cardiac and vascular anomalies as an indirect mechanism causing abnormal thyroid development.

## Methods

### Zebrafish husbandry and embryo culture

Zebrafish embryos were collected by natural spawning. Embryos were raised at 28.5°C under standard conditions (34), staged according to hours post fertilization (hpf) as described (35) and enzymatically dechorionated by incubation in embryo medium containing 0.6 mg/mL pronase (Sigma) at room temperature. For live analyses and prior to sampling and fixation, embryos were anesthetized in 0.02% tricaine (Sigma). The following zebrafish lines were used in this study: pigmentless *casper* strain (36), *Tg*(*tg:nlsEGFP*) abbreviated as *tg-nlsEGFP* (18), *Tg(sox17:EGFP)*^*s870*^ abbreviated as *sox17-EGFP* (37), *Tg(myl7:EGFP)*^*twu26*^ abbreviated as *myl7-EGFP* (38) and *Tg(kdrl:EGFP)*^*s843*^ abbreviated as *kdrl-EGFP* (39). To generate transgenic *sox17-EGFP, myl7-EGFP* and *kdrl-EGFP* embryos, homozygous transgene carriers of each line were crossed with WT fish. Reporter expression in transgenic *sox17-EGFP, myl7-EGFP* and *kdrl-EGFP* embryos highlights endodermal, myocardial and endothelial/endocardial cells, respectively. To generate pigmentless transgenic *tg-nlsEGFP* embryos, homozygous carriers of the *tg-nlsEGFP* transgene maintained in the *casper* background were crossed with non-transgenic *casper* fish. In transgenic *tg-nlsEGFP* embryos, a nuclear GFP variant is specifically expressed in *thyroglobulin*-expressing thyroid cells. If indicated, pigmentation of embryos was inhibited by addition of 0.003% 1-phenyl-2-thiourea (PTU; Sigma) to the embryo medium. Embryos were fixed in 4% phosphate-buffered paraformaldehyde (PFA; Sigma) overnight at 4°C, washed in phosphate-buffered saline containing 0.1% Tween 20 (PBST), gradually transferred to 100% methanol, and stored at −20°C until further use. Zebrafish husbandry and all experiments were performed under standard conditions in accordance with institutional (Université Libre de Bruxelles-ULB) and national ethical and animal welfare guidelines and regulation. All experimental procedures were approved by the ethical committee for animal welfare (CEBEA) from the Université Libre de Bruxelles (protocols 578N-579N).

### Preparation of Treatment Solutions

A total of 20 small molecule compounds (see Table 1 and Supplementary Table 1) were used in the pharmacological screens and each compound was applied at three concentrations. Stock solutions of test compounds were prepared in dimethylsulfoxide (DMSO, Sigma) with the exception of cyclopamine (prepared in ethanol) and SAG (prepared in distilled water). Stock solutions were aliquoted and stored at −20°C until use. Concentrations of stock solutions and test solutions are provided in Supplementary Table 1. Treatment solutions were prepared by serially diluting stock solutions in embryo medium. Control treatments included a water control group (embryo medium alone), DMSO vehicle control groups (medium containing 0.1 – 1.0% DMSO) and ethanol vehicle controls (medium containing 0.1 – 0.5% ethanol). For the gastrula screen, test solutions were additionally supplemented with 0.01 mg/L methylene blue to prevent the growth of bacteria or fungus. For the pharyngula screen, PTU was added to media to prevent pigmentation.

**Table 1.** Small molecule compounds used for modulation of the indicated signaling pathways.

| Pathway | Compound | Activity | Stock solution | Solvent | Test concentration [μM] | | |
| --- | --- | --- | --- | --- | --- | --- | --- |
|  |  |  |  |  | low | mid | high |
| BMP | DMH1 | inhibitor | 10 mM | DMSO | 3.0 | 6.0 | 12 |
| BMP | LDN193189 | inhibitor | 10 mM | DMSO | 2.0 | 4.0 | 8.0 |
| FGF | SU5402 | inhibitor | 10 mM | DMSO | 5.0 | 10 | 20 |
| FGF | PD166866 | inhibitor | 20 mM | DMSO | 2.0 | 4.0 | 8.0 |
| FGF | BCI | activator | 10 mM | DMSO | 5.0 | 10 | 20 |
| Wnt | IWR-1 | inhibitor | 20 mM | DMSO | 5.0 | 10 | 20 |
| Wnt | azakenpaullone | activator | 10 mM | DMSO | 1.25 | 2.5 | 5.0 |
| Wnt | BIO | activator | 20 mM | DMSO | 1.25 | 2.5 | 5.0 |
| Nodal/TGFβ | LY364947 | inhibitor | 5.0 mM | DMSO | 10 | 20 | 40 |
| Nodal/TGFβ | SB505124 | inhibitor | 20 mM | DMSO | 10 | 20 | 40 |
| Shh | cyclopamine | inhibitor | 10 mM | Ethanol | 10 | 25 | 50 |
| Shh | purmorphamine | activator | 10 mM | DMSO | 10 | 25 | 50 |
| Shh | SAG | activator | 5.0 mM | H<sub>2</sub>O | 2.5 | 5.0 | 10 |
| PDGF | KI11502 | inhibitor | 2.0 mM | DMSO | 0.5 | 1.0 | 2.0 |
| PDGF | AG1295 | inhibitor | 20 mM | DMSO | 7.5 | 15 | 30 |
| VEGF | vatalanib | inhibitor | 2.0 mM | DMSO | 1.25 | 2.5 | 5.0 |
| VEGF | SU4312 | inhibitor | 40 mM | DMSO | 10 | 20 | 40 |
| VEGF | GS4012 | activator | 35 mM | DMSO | 1.25 | 2.5 | 5.0 |
| Notch | LY411575 | inhibitor | 10 mM | DMSO | 2.5 | 5.0 | 10 |
| Notch | RO4929097 | inhibitor | 50 mM | DMSO | 6.0 | 12 | 25 |

### Experimental Conditions of Gastrula Screen

For signaling pathway manipulation during gastrula stages (treatment period I in Fig. 1), embryos were treated with small molecule compounds from 6 to 10 hpf. Normally developing embryos were collected from different clutches and pooled at 4 hpf. Embryos were then randomly allocated into petri dishes (60 mm × 15 mm, Sarstedt) so that each dish contained either 30 *tg-nlsEGFP* embryos or a pool of ten *sox17-EGFP*, ten *myl7-EGFP* and ten *kdrl-EGFP* embryos. At 6 hpf, embryo media were removed and 6 mL of treatment solution was added per dish. All treatments were performed in duplicate. Embryos were incubated in the dark at 28.5°C until 10 hpf. Treatments were stopped by replacing treatment solutions with embryo medium followed by three 10 min washes of embryos in fresh medium. Thereafter, embryos were transferred to clean petri dishes containing 10 mL of embryo medium and were further raised under standard conditions. At 20/21 hpf, all embryos were dechorionated and transferred to medium containing PTU. At 22 hpf, all *sox17-EGFP* embryos were inspected for gross organismal effects and GFP reporter expression and then fixed in 4% PFA. At 26/27 hpf, all remaining embryos were inspected for gross organismal effects and reporter expression. At 28 hpf, half of the *tg-nlsEGFP* embryos from each dish were fixed in 4% PFA. Remaining embryos were raised in medium containing PTU until 54/55 hpf, inspected for gross organismal effects, analyzed for reporter expression and then fixed in 4% PFA for further analyses.

**Figure 1.**
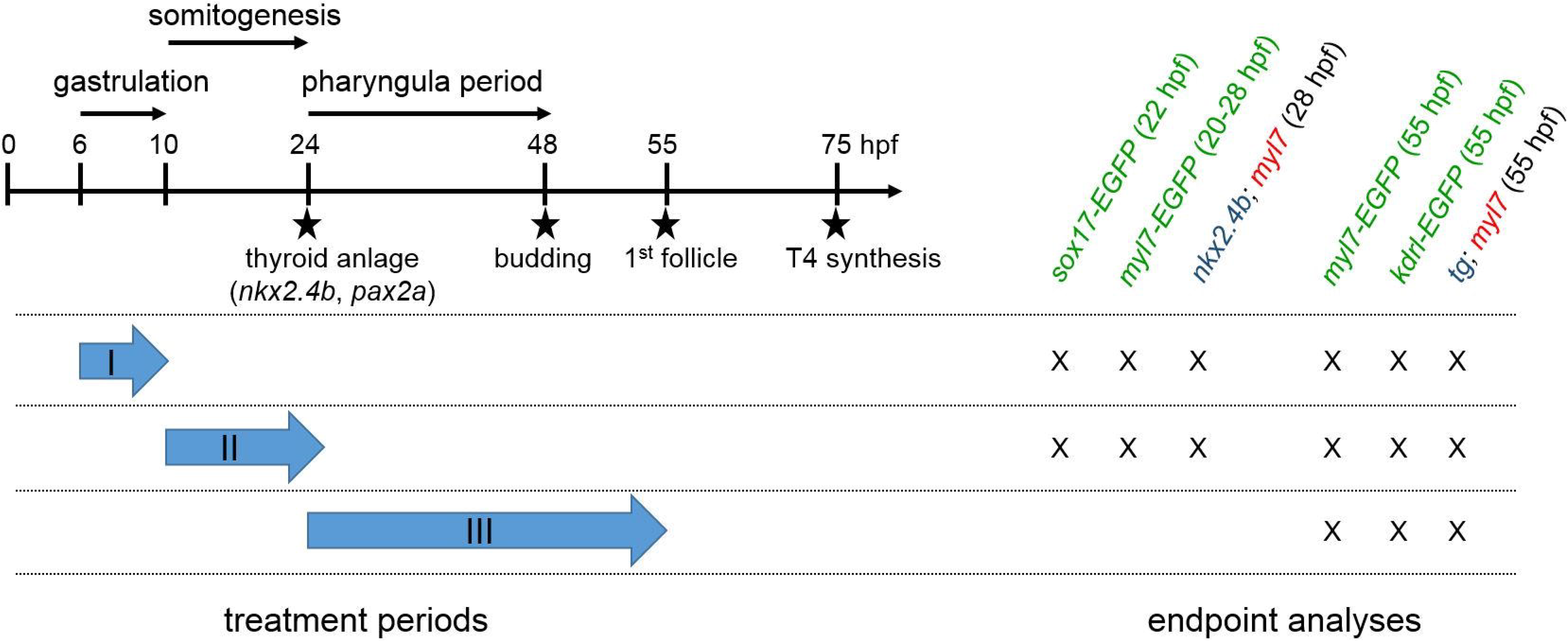
Timing of small molecule treatments and main endpoint analyses. Experimental design. The left of the schematic illustration highlights how periods of small molecule treatment during screening assays I (gastrula screen), II (somitogenesis screen) and III (pharyngula screen) relate temporally to early zebrafish development and key events of thyroid morphogenesis. Time points along the arbitrarily scaled axis are given in hours post fertilization (hpf). Note that treatment periods in the gastrula screen (6 – 10 hpf) and somitogenesis screen (10 – 26 hpf) cover developmental periods preceding thyroid anlage formation (around 23/24 hpf) whereas the treatment period in the pharyngula screen (24 – 54 hpf) covers the developmental period after the onset of thyroid anlage formation. The right part of the scheme summarizes the timing of main endpoint analyses for each assay. Endpoints include examination of thyroid (*nkx2.4b, tg*) and cardiac (*myl7*) marker expression by *in situ* hybridization and evaluation of GFP reporter expression patterns in transgenic zebrafish embryos (marked by green letters).

### Experimental Conditions of Somitogenesis Screen

Small molecule treatment during somitogenesis stages (treatment period II in Fig. 1) lasted from 10 to 26 hpf. At 8 hpf, normally developing embryos were collected from different clutches and pooled. Embryos were then randomly allocated into petri dishes (60 mm × 15 mm) as described for the gastrula screen. At 10 hpf, embryo medium was replaced by 6 mL of treatment solution per dish and embryos were incubated in test solutions at 28.5°C in the dark. All treatments were performed in duplicate. At 21 hpf, treatment of *sox17-EGFP* embryos was stopped, embryos were dechorionated, inspected for gross organismal effects and GFP reporter expression and fixed at 22 hpf in 4% PFA. All other treatments were stopped at 26 hpf by replacing treatment solutions with embryo medium followed by two 10 min washes in medium containing PTU. Embryos were dechorionated and further raised under standard conditions. At 27/28 hpf, embryos were inspected for gross organismal effects and reporter expression. At 28 hpf, half of the *tg-nlsEGFP* embryos in each dish were fixed in 4% PFA. Remaining embryos were raised in medium containing PTU to 54/55 hpf, inspected for gross organismal effects and reporter expression and fixed at 55 hpf in 4% PFA for further analyses.

### Experimental Conditions of Pharyngula Screen

Drug treatments covering pharyngula stages (treatment period III in Fig. 1) were performed from 24 to 54 hpf. For these experiments, embryos were dechorionated at 20/21 hpf, transferred to medium containing PTU and normally developing embryos were randomly allocated into petri dishes (60 × 15 mm) so that each dish contained either 20 *tg-nlsEGFP* embryos or a pool of ten *myl7-EGFP* and ten *kdrl-EGFP* embryos. At 24 hpf, embryo medium was removed and 6 mL of treatment solution was added per dish. All treatments were performed in duplicate. Embryos were incubated in treatment solutions in the dark at 28.5°C until 54 hpf. Treatments were stopped by discarding the treatment solutions followed by two 10 min washes in 10 mL of embryo medium containing PTU. All embryos were inspected for gross organismal effects and reporter expression and subsequently fixed in 4% PFA for further analyses.

### Whole-mount in situ hybridization (WISH)

DNA templates for synthesis of *nkx2.4b* (previously *nkx2.1a*, ZDB-GENE-000830-1) and *tg* (ZDB-GENE-030519-1) riboprobes were generated by PCR as described (14). The plasmid for the *myl7* (previously *cmlc2*, ZDB-GENE-991019-3) riboprobe has been used as described (40). Single color WISH of *nkx2.4b* was performed essentially as described (14) using *nkx2.4b* riboprobes labeled with digoxigenin (DIG), anti-DIG antibody conjugated to alkaline phosphatase (1:6000, Roche) and BM Purple (Roche) as alkaline phosphatase substrate.

To facilitate a higher throughput of dual-color WISH assays for *nkx2.4b*/*myl7* in 28 hpf embryos and *tg*/*myl7* in 55 hpf embryos, we developed a WISH protocol based on triple probe hybridization of pools of 28 and 55 hpf embryos using *nkx2.4b, tg* and *myl7* riboprobes labeled with DIG, dinitrophenol (DNP) and fluorescein (FLU), respectively. We first confirmed for each experimental treatment group that embryos collected at 28 and 55 hpf can be unequivocally distinguished by morphological landmarks so that these embryos can be pooled during different steps of WISH assay. We additionally identified many experimental treatment groups displaying unique, treatment-related morphological anomalies (i.e. cyclopia, curled tails, etc.) that would permit pooling with normally developing embryos from other treatments.

Embryos were processed during pre-hybridization protocol steps as described (41). Just prior to probe hybridization, stage 28 and 55 hpf embryos were pooled in hybridization tubes according to morphological criteria and incubated in 1 mL triple-probe hybridization buffer for at least 16 hours at 65°C. Post-hybridization washes were performed with pooled samples as described (41). Before antibody incubation, embryos were sorted for developmental stage. Stage 28 hpf embryos were incubated with anti-DIG to detect the *nkx2.4b* probe and stage 55 hpf embryos were incubated with anti-DNP antibody conjugated to alkaline phosphatase (1:500, Vector Laboratories) to detect *tg* probe. After overnight antibody incubation and several washing steps, *nkx2.4b* expression in 28 hpf embryos was revealed by BM purple staining and *tg* expression in 55 hpf embryos by NBT/BCIP staining (Roche). Staining reactions were stopped by several 5 min washes in PBST followed by a 10 min incubation in 100% methanol to remove background staining. Thereafter, stage 28 and 55 hpf embryos were recombined for two 5 min HCl-glycine (pH 2.2) treatments to remove the first antibody (42) followed by overnight incubation of pooled samples with anti-FLU antibody conjugated to alkaline phosphatase (1:2000, Roche). Expression of *myl7* in 28 and 55 hpf embryos was then revealed by Fast Red (Sigma) staining. Double-stained embryos were washed in PBST, post-fixed in 4% PFA and embedded in 90% glycerol. Stained specimen were sorted according to developmental stage and treatment group and analyzed for treatment-related effects on thyroid and cardiac marker expression. Whole-mount imaging of stained specimen was performed using a DFC420C digital camera mounted on a MZ16F stereomicroscope (Leica). If indicated, embryos were embedded in 7% low-melting point agarose (Lonza) and sagittal tissue sections at 100 µm thickness were cut on a VT1000S vibratome (Leica). Sections were mounted in Glycergel (Dako) and images were acquired using an Axiocam digital camera mounted on an Axioplan 2 microscope (Zeiss).

### Whole-mount immunofluorescence (WIF)

WIF of transgenic embryos expressing EGFP in endoderm, myocardium or endothelium/endocardium was performed essentially as described (14). EGFP expression was detected using chicken anti-GFP polyclonal antibody (1:1000; Abcam, ab13970) and Alexa Fluor 488-conjugated goat anti-chicken IgG antibody (1:250; Invitrogen, A-11039). Stained specimens were post-fixed in 4% PFA, gradually transferred to 90% glycerol and phenotypically analyzed using a M165 FC fluorescence stereomicroscope. Whole mount imaging of stained specimens was performed with a DMI600B epifluorescence microscope equipped with a DFC365FX camera (Leica). If indicated, embryos were embedded in 1% low-melting point agarose on Fluoro-Dish glass bottom dishes (World Precision Instruments) and whole mount confocal imaging was performed with a LSM 510 confocal microscope (Zeiss) using Zen 2010 D software (Zeiss).

## Results

### Pharmacological treatments and developmental endpoint analyses

Using a pharmacological approach, our study aimed at temporally dissecting the role of major signaling pathways for normal thyroid morphogenesis in zebrafish embryos. The small molecules used in study (see Table 1) were selected based on a thorough review of available zebrafish studies reporting on their efficacy and specificity to modulate the activity of specific signaling pathways. For all compounds, maximum tolerable concentrations were determined during initial range-finding studies to ensure low mortality rates during the definitive treatment experiments.

Limited solubility in DMSO or ethanol prevented the preparation of highly concentrated stock solutions for some compounds (Table 1). As a consequence, some treatment solutions contained increased solvent concentrations of up to 1% DMSO or 0.5% ethanol. To account for the different solvent concentrations used during the definitive treatment studies, all experiments included three DMSO-containing vehicle control groups (0.1, 0.5, 1.0%) and at least two ethanol-containing vehicle control groups (0.1, 0.5%).

Possible effects of small molecule treatment on early thyroid development were analyzed at two developmental stages. At 28 hpf, all embryos were examined for *nkx2.4b* mRNA expression by means of WISH. While first *nkx2.4b*-expressing TPC can be identified in zebrafish as early as 23/24 hpf, a larger domain of thyroidal *nkx2.4b* expression with stereotypic dimension and localization can be visualized by WISH in 28 hpf embryos (see embryo #3 in Fig. 2). We refer to this expression domain as the zebrafish thyroid anlage. The absence of other tissues expressing *nkx2.4b* in the vicinity of the thyroid anlage facilitates sensitive analyses to identify treatment-related changes in size, shape and localization of the thyroid anlage. Within the anterior endoderm, the thyroid anlage normally forms in a region adjacent to the developing cardiac OFT (see embryo #1 in Fig. 2). Co-staining of 28 hpf embryos with a probe for the myocardial marker *myl7* provided also relevant landmark information to identify possible ectopic locations of *nkx2.4b* expression. In addition to topological properties of the thyroid anlage, we carefully monitored the progressive evolution of the staining signal in all WISH assays as an indicator for possible treatment-related differences in *nkx2.4b* expression levels. If robustly detected across replicate experiments, we will refer to such observations as enhanced or diminished expression levels.

**Figure 2.**
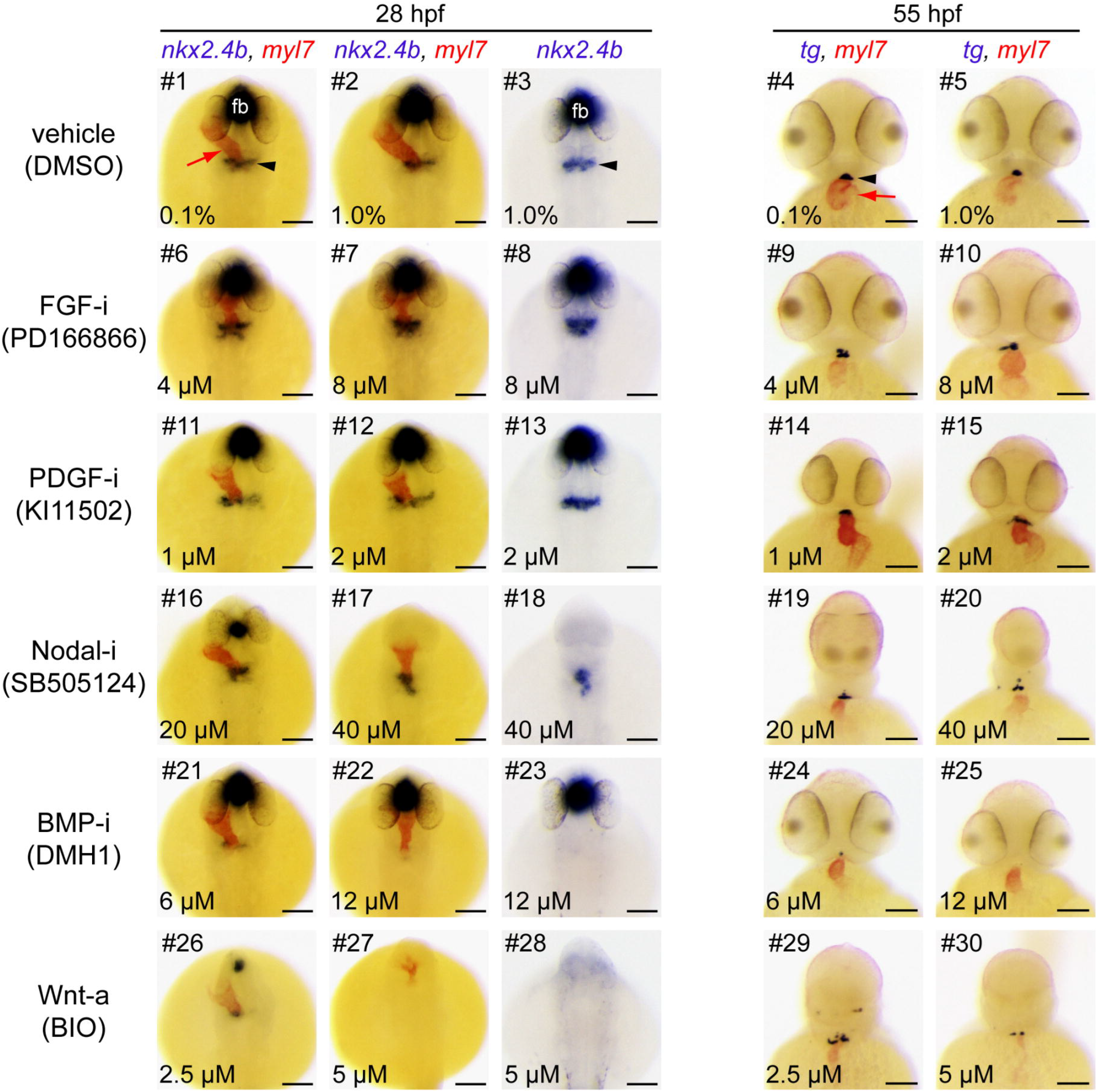
Major thyroid phenotypes recovered in the gastrula screen. Zebrafish embryos were treated from 6 to 10 hpf with inhibitors (i) or activators (a) of major signaling pathways. At 28 and 55 hpf, expression of thyroid (*nkx2.4b, tg*) and cardiac (*myl7*) markers was assessed by single and dual-color *in situ* hybridization. Note that *nkx2.4b* is expressed in the thyroid anlage and the ventral forebrain (fb). Black arrowheads point to domains of thyroid marker expression and red arrows point to the developing heart. Phenotypic comparison of thyroid development in treated and control embryos (DMSO, dimethylsulfoxide) at 28 hpf revealed enhanced expression of the early thyroid marker *nkx2.4b* (#8,13), aberrant posterior expansion of the thyroidal *nkx2.4b* expression domain (#6,8,17,18) and severe reduction or complete absence of thyroidal *nkx2.4b* expression (#21-23, #26-28). Analysis of *tg* expression in 55 hpf embryos showed irregular morphologies of the thyroid primordium (#9,10,15,19,20,29), enlarged thyroids (#15), smaller thyroids (#20,24,30) or complete absence of *tg* expression (#25). See text for more details on treatment-dependent thyroid phenotypes. Dorsal (28 hpf) and ventral (55 hpf) views are shown, anterior is to the top. Scale bars: 100 µm.

Treatment-related effects on thyroid morphogenesis were also analyzed at 55 hpf. At this developmental stage, we examined the expression characteristics of the functional differentiation marker *thyroglobulin (tg)* which was assessed by means of monitoring nlsEGFP reporter expression in live transgenic *tg-nlsEGFP* embryos (18) and by WISH analyses of endogenous *tg* mRNA expression. In 55 hpf embryos, the thyroid primordium has completed its detachment from the ventral pharyngeal endoderm and presents as a compact midline mass of thyroid cells with a slightly ovoid shape when viewed frontally (see embryo #4 in Fig. 2). All embryos were assessed for treatment-related differences in thyroid size, shape, localization and *myl7* co-staining was performed to identify possible ectopic locations of *tg* expression relative to the cardiac OFT. In addition to the evaluation of thyroid markers, drugged embryos were assessed for treatment-related gross morphological anomalies (edema, tissue malformations, developmental delay, growth deficits) by brightfield microscopy. Fluorescence microscopy of transgenic embryos was used to qualitatively assess possible deviations from normal endoderm (*sox17-EGFP* line), cardiac (*myl7-EGFP* line) and vascular (*kdrl-EGFP* line) development.

### Pathway Manipulation during Gastrula Stages

A first pharmacological screen addressed thyroid development following manipulation of signaling pathways during gastrula stages. This treatment (6 to 10 hpf) coincides with the developmental period during which the three germ layers (ectoderm, mesoderm, endoderm) and the major body axes are formed. Many of the pathways targeted in our experiments have critical regulatory roles during these processes, most notably BMP, FGF, Wnt and Nodal pathways (43, 44). Accordingly, the presence of characteristic developmental anomalies in drugged embryos allowed us to verify the efficacy of pathway disruption. Small molecule-induced abnormalities included C2 dorsalized phenotypes after BMP inhibition (45), loss of caudal tissue in FGF inhibitor-treated embryos (Supplementary Fig. 2A) (46), cyclopic eyes and loss of forebrain following Nodal inhibitor treatment (47) and neural posteriorization phenotypes in embryos treated with Wnt activators (48).

#### Thyroidal Effects

Small molecule treatment of zebrafish embryos covering gastrula stages revealed alterations in early thyroid development in response to nine drugs affecting five different pathways. Specifically, abnormal thyroid development resulted from treatment with inhibitors of FGF, platelet-derived growth factor (PDGF), Nodal/transforming growth factor β (TGFβ) and BMP pathways and activators of Wnt signaling (Fig. 2,3).

**Figure 3.**
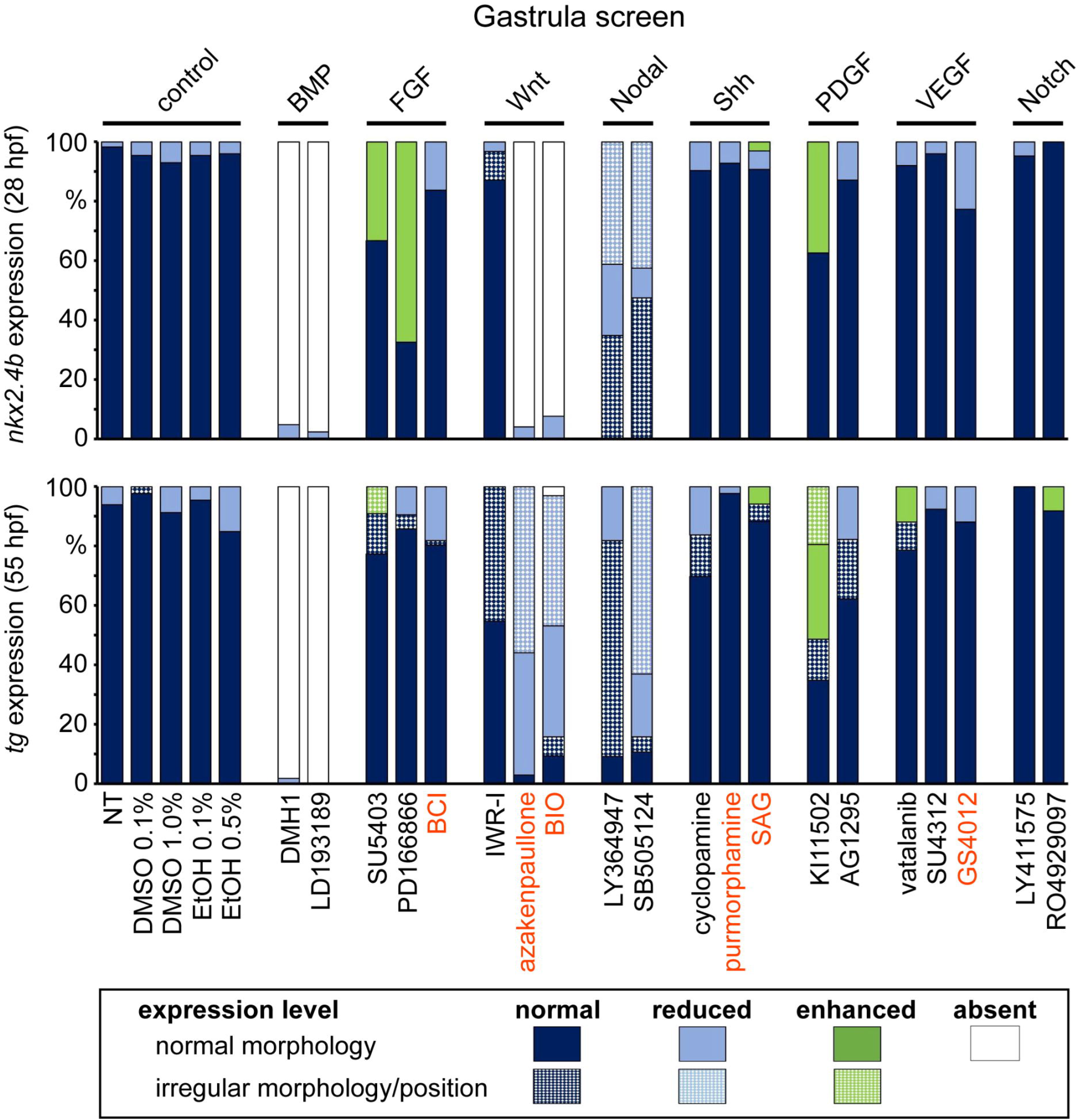
Distribution of thyroid phenotypes recovered in the gastrula screen. Zebrafish embryos were treated from 6 to 10 hpf with a panel of 20 small molecule compounds known to inhibit (black letters) or activate (red letters) specific signaling pathways. Control groups included non-treated embryos (NT) and embryos treated with dimethylsulfoxide (DMSO) or ethanol (EtOH). Bar graphs depict quantification of the proportion of specimen displaying abnormal thyroid development as determined by *nkx2.4b* staining of 28 hpf embryos (upper panel) and *tg* staining of 55 hpf embryos (lower panel). Results are shown for the highest test concentration of each compound and data are presented as the percentage of embryos displaying a particular phenotype. Thyroid phenotypes were classified into seven categories according to the overall expression level of the marker gene and apparent deviations from normal morphology and/or positioning of the expression domain (highlighted by texture overlays).

At 28 hpf, embryos treated with two different FGF pathway inhibitors (PD166866, SU5402) showed a more rapid evolution of the staining for the early thyroid marker *nkx2.4b* compared to vehicle-treated embryos (Fig. 2). In addition, we frequently detected a posterior expansion of the thyroidal *nkx2.4b* expression domain (see embryo #8 in Fig. 2) or additional ectopic domains of *nkx2.4b* expression (Supplementary Fig. 1). Analysis of *sox17-EGFP* embryos did not reveal discernible effects of FGF inhibition on anterior endoderm formation (Fig. 4). Notably, comparative analyses of *tg* staining between drugged embryos and control embryos did not reveal gross differences in thyroid size at 55 hpf (Fig. 2). However, FGF inhibitor-treated embryos presented at 55 hpf a less compact mass of thyroid cells (see embryo #9 in Fig. 2) and irregular thyroid morphology (see embryo #10 in Fig. 2 and Supplementary Fig. 2B). Conversely, treatment with BCI, a dusp6 inhibitor that locally enhances FGF signaling, did not cause clear thyroid effects as small reductions in thyroid size were limited to embryos displaying an overall developmental delay.

**Figure 4.**
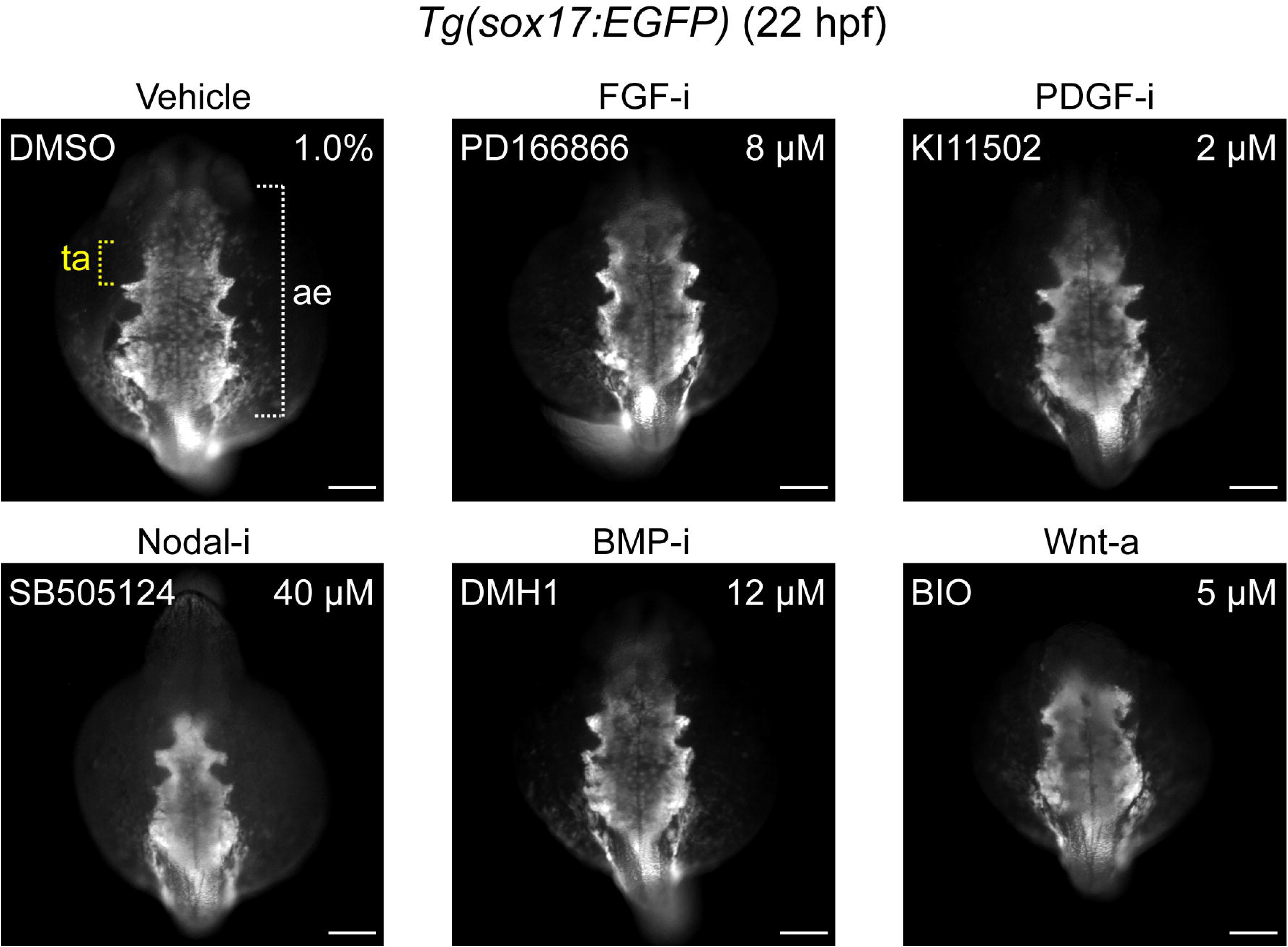
Effects of small molecule treatment during gastrulation on anterior endoderm formation. Transgenic *Tg(sox17:EGFP)* embryos were treated from 6 to 10 hpf with inhibitors (i) or activators (a) of specific signaling pathways and GFP reporter expression was analyzed by immunofluorescence staining at 22 hpf to assess formation of the anterior endoderm (ae). The prospective region of thyroid anlage (ta) formation within the anterior endoderm is highlighted by yellow brackets. Phenotypic comparison between treated and control embryos (DMSO, dimethylsulfoxide) revealed a severe reduction of anterior endoderm in embryos treated with inhibitors of Nodal signaling and perturbed morphogenesis of lateral and most rostral endoderm following overactivation of Wnt signaling. Dorsal views are shown, anterior is to the top. Scale bars: 100 µm.

Enhanced *nkx2.4b* expression at the thyroid anlage stage was observed also in embryos treated with KI11502, a receptor tyrosine kinase inhibitor with selectivity for PDGF signaling (Fig. 2). A modest mediolateral expansion of the thyroidal *nkx2.4b* expression domain was observed in approximately one third of KI11502-treated embryos (see embryo #13 in Fig. 2). Endoderm formation was unaffected by Ki11502 treatment (Fig. 4). Thyroid anomalies due to Ki11502 treatment were also apparent at later stages including frequent detection of an irregular mediolateral expansion of *tg* expression in 55 hpf embryos (see embryo #15 in Fig. 2). The increased thyroid size in KI11502-treated embryos was particularly remarkable given that these embryos displayed reduced whole body and head size at 55 hpf (see embryo #15 in Fig. 2).

Consistent with the crucial role of Nodal signaling in early embryonic development, inhibitors of this pathway (LY364947 and SB505124) caused, in a concentration-dependent manner, severe gross morphological defects including loss of ventral forebrain structures (see lack of forebrain *nkx2.4b* expression in embryos #17,18 in Fig. 2), cyclopia (see embryos # 19,20 in Fig. 2) and a dramatic reduction of anterior endoderm (Fig. 4). Despite these overt developmental anomalies, thyroidal *nkx2.4b* expression was robustly detectable in 28 hpf embryos, even in embryos completely lacking forebrain *nkx2.4b* expression (see embryos #17,18 in Fig. 2). However, shape and position of the thyroid anlage was abnormal in most inhibitor-treated embryos. Specifically, the thyroidal domain of *nkx2.4b* expression showed a reduced mediolateral extension but expanded further caudally (see embryo #18 in Fig. 2). In embryos treated with high concentrations of inhibitors, additional ectopic clusters of *nkx2.4b*-expressing cells were frequently detected in posterior positions (Supplementary Fig. 3). At 55 hpf, the thyroid primordium of most embryos from LY364947 and SB505124 treatments displayed an irregular shape and the number of *tg*-expressing cells appeared reduced in SB505124-treated embryos if compared to controls (see embryo #20 in Fig. 2).

The most severe defects in early thyroid development resulted from inhibition of BMP signaling and from over-activation of canonical Wnt signaling. Treatment of gastrulating embryos with two BMP pathway inhibitors (DMH1 and LDN193189) consistently caused a concentration-dependent loss of thyroidal *nkx2.4b* expression at 28 hpf. Almost all embryos treated with the highest inhibitor concentrations displayed a complete lack of detectable *nkx2.4b* expression in the thyroid region. The absence of a thyroid anlage was not associated with any discernible reduction of foregut endoderm formation as judged from analyses of *sox17-EGFP* embryos (Fig. 4). For both BMP pathway inhibitors, early defects in thyroid specification correlated closely with a severe reduction or absence of detectable *tg* staining in embryos at later stages (see embryos #24,25 in Fig. 2).

Over-activation of canonical Wnt signaling in gastrulating embryos by treatment with BIO and azakenpaullone caused complex developmental phenotypes. At 28 hpf, drugged embryos showed a concentration-dependent loss of anterior neural tissue concurrent with severe inhibition of cardiac development and a dramatic reduction of thyroidal *nkx2.4b* expression (see embryos #26-28 in Fig. 2). At the highest test concentrations of BIO (5.0 µM) and azakenpaullone (5.0 µM), *nkx2.4b* expression was no longer detectable in the thyroid region. Although formation of anterior endoderm was evident in embryos treated with Wnt activators, perturbed morphogenesis was frequently observed in lateral and most rostral regions of the anterior endoderm (Fig. 4). Interestingly, despite the absence of detectable *nkx2.4b* expression in the majority of 28 hpf embryos treated with high drug concentrations, all 55 hpf embryos had detectable *tg* expression, although *tg* expression was restricted to tiny and often ectopically positioned domains (see embryo #30 in Fig. 2). Conversely, inhibition of Wnt signaling by IWR1 treatment did not affect thyroid specification. Deviations from normal thyroid development were limited to an aberrant shape of the thyroid primordium in 55 hpf embryos when viewed laterally (Supplementary Fig. 2B).

### Pathway Manipulation during Somitogenesis Stages

A second pharmacological screen involved treatment of embryos from 10 to 26 hpf and covered the somitogenesis period of zebrafish embryonic development. In zebrafish, the endoderm becomes morphologically distinctive at the beginning of the somitogenesis period as two converging sheets of cells that will fuse at the midline by the 15-17 somite stage (49). While the posterior endoderm rapidly remodels into a rod-like structure, the prospective foregut endoderm region will maintain a sheet-like morphology until after completion of the somitogenesis. Towards the end of the somitogenesis period, TPC co-expressing *nkx2.4b, pax2a* and *hhex* become detectable around 23/24 hpf in the anterior endoderm region (31).

#### Thyroidal effects

Somitogenesis treatment of zebrafish embryos with small molecules revealed alterations of early thyroid development in response to 11 drugs affecting seven different pathways (Fig. 5 and 6). Drug-induced deviations from normal thyroid development were detectable following treatment with inhibitors of PDGF, vascular endothelial growth factor (VEGF), Sonic hedgehog (Shh), Nodal/TGFβ, BMP and FGF signaling and activators of Wnt signaling (Fig. 7).

**Figure 5.**
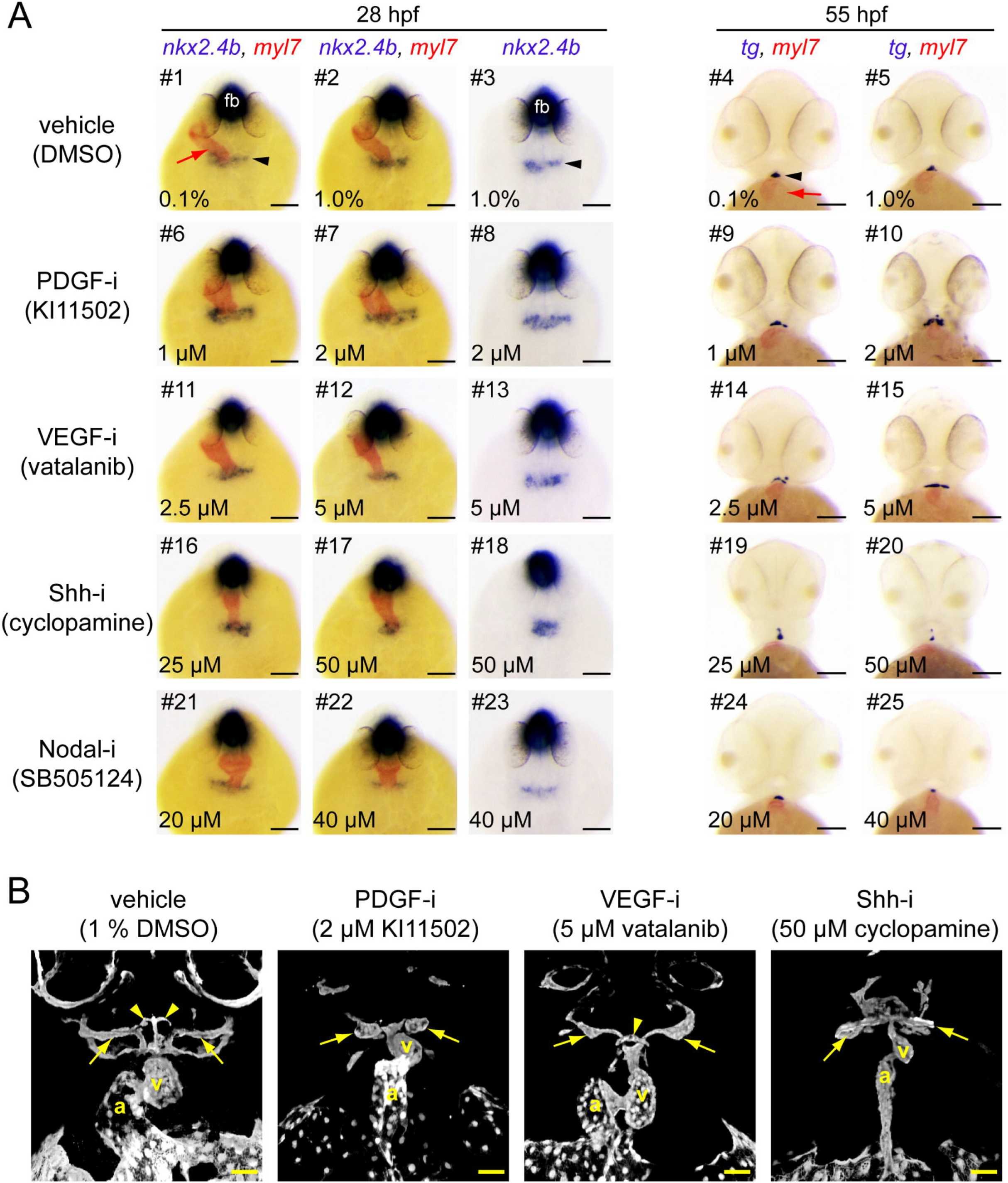
Thyroid and vascular phenotypes recovered in the somitogenesis screen. **(A)** Embryos treated from 10 to 26 hpf with inhibitors (i) of PDGF, VEGF, Shh and Nodal signaling pathways were analysed by single and dual-color *in situ* hybridization for expression of thyroid (*nkx2.4b, tg*) and cardiac (*myl7*) markers. Note that *nkx2.4b* is expressed in the thyroid anlage and the ventral forebrain (fb). Black arrowheads point to domains of thyroid marker expression and red arrows point to the developing heart. Phenotypic comparison of thyroid development in treated and control embryos (DMSO, dimethylsulfoxide) at 28 hpf revealed a mediolateral expansion (PDGF-i, #6-8) or mediolateral reduction of the thyroidal *nkx2.4b* expression domain (Shh-i, #17-18) as well as diminished levels of *nkx2.4b* expression (Nodal-i, #22,23). Analysis of *tg* expression in 55 hpf embryos showed irregularly expanded domains of *tg* expression (PDGF-i, #9,10; VEGF-i, #15), irregularly positioned and reduced domains of *tg* expression (Shh-i, #20) and overtly smaller thyroids (Nodal-i, #25). See text for more details on treatment-dependent thyroid phenotypes. Dorsal (28 hpf) and ventral (55 hpf) views are shown, anterior is to the top. Scale bars: 100 µm. **(B)** Transgenic *Tg(kdrl:EGFP)* embryos were treated from 10 to 26 hpf with small molecule compounds and GFP reporter expression was analyzed by immunofluorescence staining at 55 hpf to assess morphogenesis of the pharyngeal vasculature. Confocal analysis of vascular development in control embryos (DMSO, dimethylsulfoxide) and embryos treated with inhibitors (i) of PDGF, VEGF and Shh signaling revealed treatment-induced dysplasia of major vessels proposed to guide thyroid localization. Highlighted are defects in formation of the hypobranchial artery (arrowheads) and the first aortic arch arteries (arrows). 3D reconstructions of confocal images are shown, ventral view, anterior is to the top. Abbreviations: a atrium; v ventricle. Scale bars: 50 µm.

**Figure 6.**
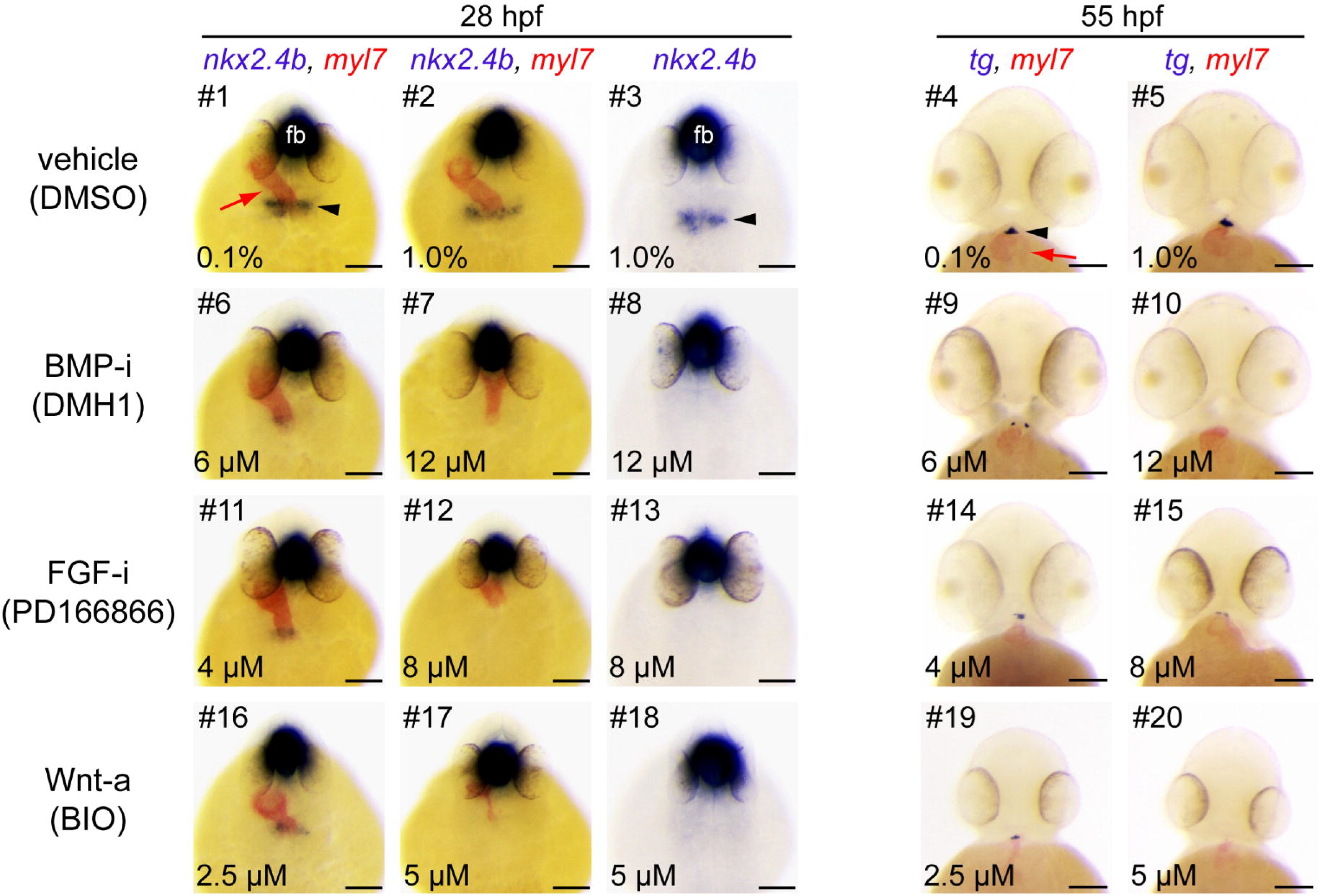
Manipulation of BMP, FGF and Wnt signaling during somitogenesis results in thyroid agenesis. Embryos treated from 10 to 26 hpf with inhibitors (i) of BMP and FGF or activators (a) of Wnt signaling pathways were analyzed by single and dual-color *in situ* hybridization for expression of thyroid (*nkx2.4b, tg*) and cardiac (*myl7*) markers. Note that *nkx2.4b* is expressed in the thyroid anlage and the ventral forebrain (fb). Black arrowheads point to domains of thyroid marker expression and red arrows point to the developing heart. Phenotypic comparison of thyroid specification in treated and control embryos (DMSO, dimethylsulfoxide) at 28 hpf revealed severe reduction or complete absence of thyroidal *nkx2.4b* expression after treatment with BMP inhibitors (BMP-i, #6-8), FGF inhibitors (FGF-i, #11-13) and activators of Wnt signaling (Wnt-a, #16-18). Analysis of 55 hpf embryos similarly showed severe reduction or complete absence of *tg* expression in response to these treatments (BMP-i, #9,10; FGF-i, #14,15; Wnt-a, #19,20). See text for more details on treatment-dependent thyroid phenotypes. Dorsal (28 hpf) and ventral (55 hpf) views are shown, anterior is to the top. Scale bars: 100 µm.

**Figure 7.**
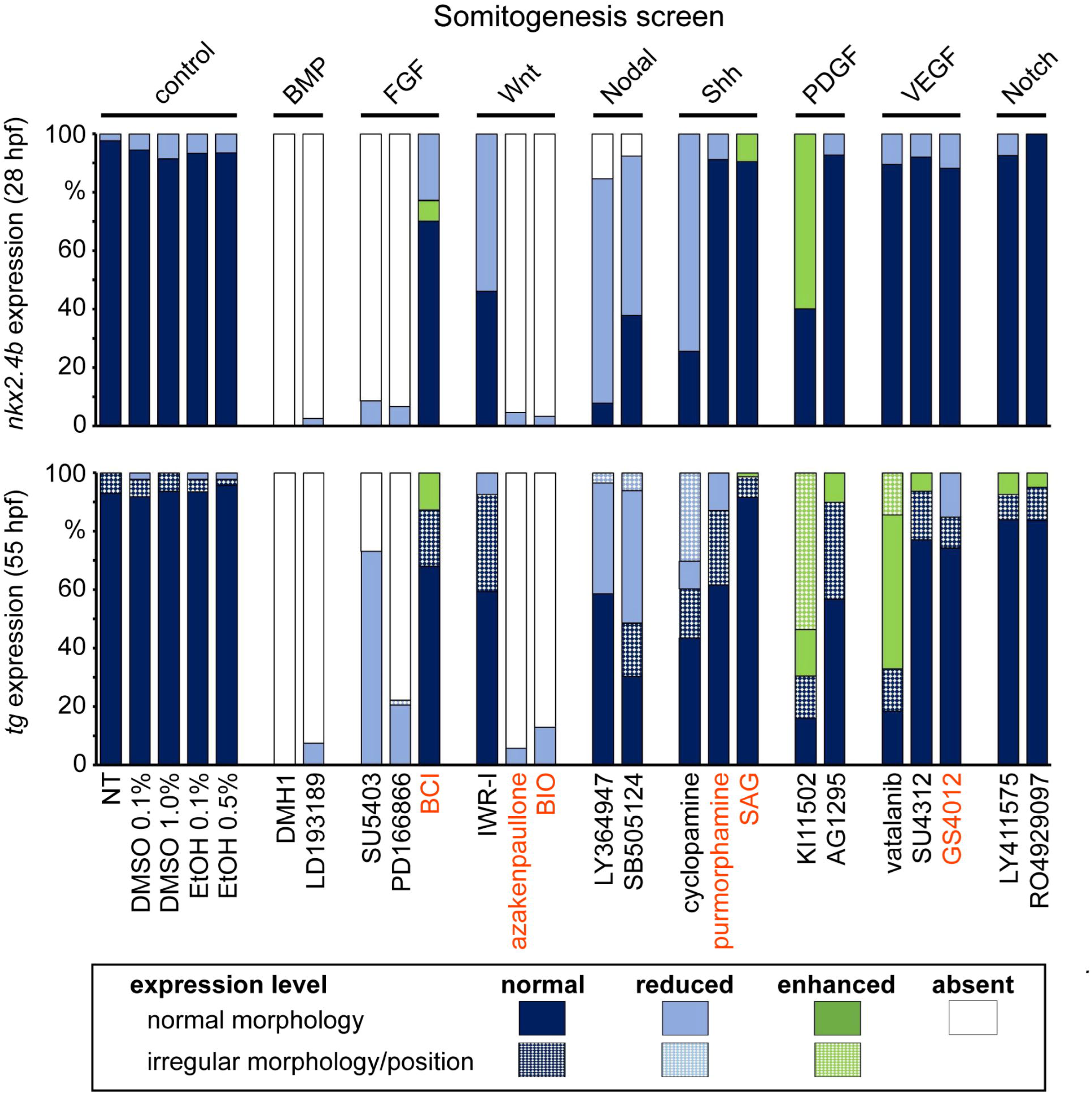
Distribution of thyroid phenotypes recovered in the somitogenesis screen. Zebrafish embryos were treated from 10 to 26 hpf with a panel of 20 small molecule compounds known to inhibit (black letters) or activate (red letters) specific signaling pathways. Control groups included non-treated embryos (NT) and embryos treated with dimethylsulfoxide (DMSO) or ethanol (EtOH). Bar graphs depict quantification of the proportion of specimen displaying abnormal thyroid development as determined by *nkx2.4b* staining of 28 hpf embryos (upper panel) and *tg* staining of 55 hpf embryos (lower panel). Results are shown for the highest test concentration of each compound and data are presented as the percentage of embryos displaying a particular phenotype. Thyroid phenotypes were classified into seven categories according to the overall expression level of the marker gene and apparent deviations from normal morphology and/or positioning of the expression domain (highlighted by texture overlays).

Inhibition of PDGF signaling during somitogenesis by KI11502 treatment caused a marked mediolateral expansion of the *nkx2.4b* expression domain at 28 hpf (see embryos #6-8 in Fig. 5A). This enlargement of the thyroid anlage occurred concomitantly with an abnormal heart morphology characterized by a widened arterial pole region of an otherwise shorter heart tube (see embryo #7 in Fig. 5A). While endoderm formation was unaffected by PDGF inhibition, live analyses of heart development in inhibitor-treated *myl7-EGFP* embryos revealed delayed cardiac fusion and heart tube assembly (Supplementary Fig. 4) as previously described in zebrafish *pdgfra* mutants (50). At 55 hpf, the thyroid primordium of KI11502-treated embryos appeared enlarged compared to control embryos. The expanded domains of *tg* expression in drugged embryos had an aberrant morphology often characterized by an irregular bilateral expansion (see embryos #9,10 in Fig. 5A).

A very similar late thyroid phenotype was observed in embryos following treatment with the VEGF inhibitor vatalanib (see embryo #15 in Fig. 5A). Concurrent analyses of pharyngeal vasculature development in *kdrl-EGFP* embryos revealed another commonality between KI11502 and vatalanib treatments in that irregularly enlarged thyroids were predominantly observed in treatment groups with defective OFT vessel morphogenesis (Fig. 5B). Three-dimensional reconstructions of confocal images of the pharyngeal vasculature in KI11502- and vatalanib-treated embryos showed dysplasia of specific vessels previously identified as vital structures guiding the localization of zebrafish thyroid tissue (15).

Inhibition of Shh signaling during somitogenesis by treatment with the Smoothened antagonist cyclopamine resulted in complex phenotypic alterations. Cyclopamine treatment between 10 to 26 hpf efficiently phenocopied many malformations previously described in zebrafish mutants with impaired Shh signaling including ‘curly tail down’ body shape, mild-to-severe cyclopia, hemorrhage as well as OFT malformations (51, 52). Despite these severe developmental defects, cyclopamine treatment caused only mild alterations of the thyroid anlage in 28 hpf embryos (Fig. 5A). While the intensity of *nkx2.4b* staining in cyclopamine-treated embryos was similar to control levels, a unique topological characteristic of the thyroid anlage in cyclopamine-treated embryos was that TPC were restricted to more medial positions so that the anlage had an overall reduced mediolateral size (see embryo #17 in Fig. 5A). Prevalence of this early thyroid phenotype in cyclopamine treatments was correlated with concentration-dependent decrements in the mediolateral size of the anterior endoderm (Supplementary Fig. 5). At 55 hpf, cyclopamine-treated embryos displayed severe malformations of pharyngeal development. Embedded in a grossly abnormal subpharyngeal environment, multiple clusters of *tg*-expressing cells were detectable at ectopic locations along the pharyngeal midline (Fig. 5A). Additional confocal analyses of the pharyngeal vasculature confirmed dysgenesis of major OFT vessels in cyclopamine-treated *kdrl-EGFP* embryos (Fig. 5B).

Somitogenesis treatment with inhibitors of Nodal/TGFβ signaling (LY364947, SB505124) caused moderate thyroid phenotypes (Fig. 5A). At the thyroid anlage stage, the major phenotype was a reduced intensity of *nkx2.4b* staining while size, shape and location of the *nkx2.4b* expression domain was similar to controls (see embryo #23 in Fig. 5A). In later stage embryos, inhibition of Nodal/TGFβ signaling resulted in a concentration-dependent decrease of the size of the thyroid primordium (see embryo #25 in Fig. 5A).

In the somitogenesis screen, the most severe thyroid defects were observed following treatment of embryos with inhibitors of BMP (DMH1 and LDN193189) and FGF signaling (PD166866, SU5402) and compounds stimulating canonical Wnt signaling (BIO, azakenpaullone). For all six compounds, a concentration-dependent loss of thyroidal *nkx2.4b* expression was evident in 28 hpf embryos (Fig. 6). At the highest test concentration, each of the compounds caused a complete lack of detectable *nkx2.4b* expression in the thyroid region. Conversely, drug treatment did not cause gross defects in anterior endoderm formation as judged from analyses of *sox17-EGFP* embryos (data not shown). However, dual-color WISH of *nkx2.4b* and *myl7* expression showed that the concentration-dependent loss of thyroid marker expression caused by BIO and azakenpaullone treatment was accompanied by concurrent depletion of myocardial cells and failure of heart tube formation at high test concentrations (see embryo #17 in Fig. 6). Although inhibition of BMP and FGF signaling resulted also in heart malformations (i.e. jogging and looping defects) in a high proportion of embryos, there was no similar dramatic loss of cardiac tissue. For all treatments, impaired TPC specification in 28 hpf embryos was associated with a dramatic diminution or even complete absence of *tg*-expressing cells in 55 hpf embryos (Fig. 6). Yet, there were notable differences among differentially treated embryos with respect to the phenotypic presentation of the small remaining thyroid tissue. The small thyroid primordium of 55 hpf embryos treated with activators of Wnt signaling was mostly restricted to the expected midline position (see embryo #19 in Fig. 6). In embryos treated with BMP inhibitors, *tg*-expressing cells were frequently detected off the midline, predominantly in tiny bilateral cell clusters located on either side of the cardiac OFT (see embryo #9 in Fig. 6). Conversely, inhibition of FGF signaling caused a much more variable localization of *tg*-expressing cells with irregularly positioned clusters of thyroid cells expanding uni- or bilaterally away from the midline (see embryo #15 in Fig. 6).

### Pathway Manipulation during Pharyngula Stages

In a third screen, embryos were treated with small molecules from 24 to 54/55 hpf covering the pharyngula period of zebrafish embryogenesis. During this period, the endoderm undergoes further dramatic remodeling to finally form a primitive gut tube (53). The thyroid anlage which is formed during early pharyngula stages transforms into a thyroid bud during the second half of the pharyngula period. Notably, thyroid bud formation in zebrafish is accompanied by the onset of expression of functional differentiation markers including *tg, tshr, tpo* and *slc5a5* between 34 and 42 hpf (14). This thyroid primordium detaches from the then formed ventral floor of the foregut around 48 hpf and relocates into the subpharyngeal mesenchyme in close apposition to the cardiac OFT. According to these timelines, this screen specifically addressed the role of signaling cues for thyroid budding, bud detachment and relocation of the thyroid diverticulum.

#### Thyroidal effects

Small molecule treatment of zebrafish embryos from 24 to 54/55 hpf revealed alterations of thyroid development in response to 10 drugs affecting five six different pathways (Fig. 8,9). In this screen, embryos were analyzed only at 55 hpf and drug-induced thyroid anomalies were detectable after treatment with inhibitors of BMP, FGF, Shh, Nodal/TGFβ, PDGF and VEGF signaling.

**Figure 8.**
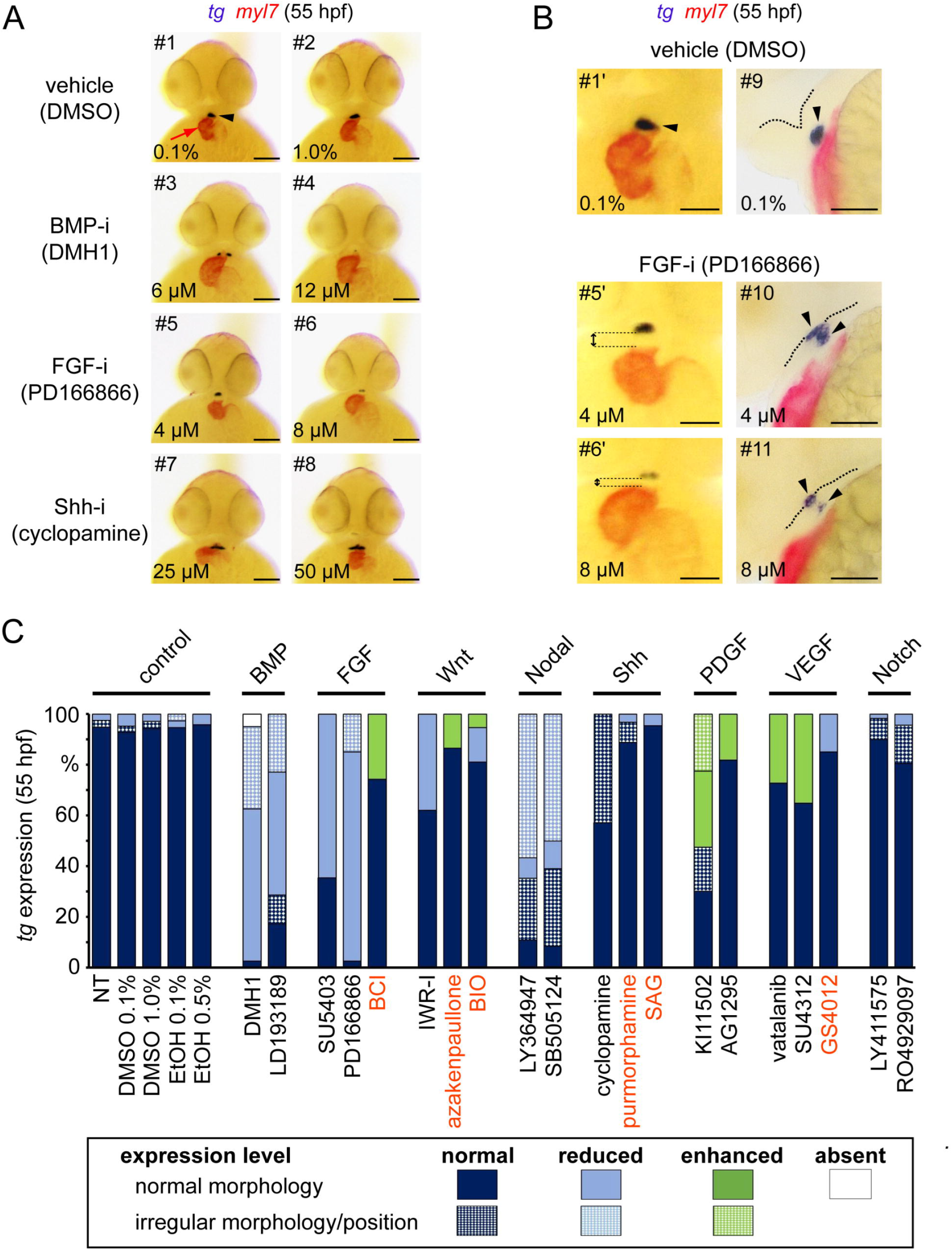
Thyroid phenotypes recovered in the pharyngula screen. **(A)** Embryos treated with small molecules from 24 to 54 hpf were analyzed at stage 55 hpf by dual-color *in situ* hybridization for thyroidal *tg* (arrowheads) and cardiac *myl7* (red arrows) expression. Phenotypic comparison of treated and control embryos (DMSO, dimethylsulfoxide) revealed reduced thyroid size after treatment with BMP inhibitors (BMP-i, #3,4) and FGF inhibitors (FGF-i, #5,6) and irregular bilateral thyroid expansion after treatment with Shh inhibitors (Shh-i, #7,8). Ventral views are shown, anterior is to the top. Scale bars: 100 µm. **(B)** The left panel of images (#1’,5’,6’) in **B** shows magnified ventral views of the cardiac outflow tract region for embryos shown in **A**. Note the apparent gap between thyroid marker expression (arrowheads) and the distal portion of the *myl7*-expressing cardiac outflow tract as observed in several embryos treated with FGF inhibitors (FGF-i, #5,6). In the right panel of images, mid-sagittal vibratome sections (anterior to the left) of stained embryos from the same experiment show that inhibition of FGF signaling results in aberrant thyroid detachment from the pharyngeal floor (indicated as dotted line). Scale bars: 50 µm. **(C)** Bar graph depicts quantification of the proportion of embryos displaying abnormal *tg* staining at stage 55 hpf in the pharyngula screen. Results are shown for non-treated embryos (NT), vehicle control embryos treated with dimethylsulfoxide (DMSO) or ethanol (EtOH) and for the highest test concentration of each compound. Data are presented as the percentage of embryos displaying a particular phenotype. Compounds with activating activity on specific signaling pathways are highlighted in red letters. Thyroid phenotypes were classified into seven categories according to the overall *tg* expression level and apparent deviations from normal morphology and/or positioning of the expression domain (highlighted by texture overlays).

**Figure 9.**
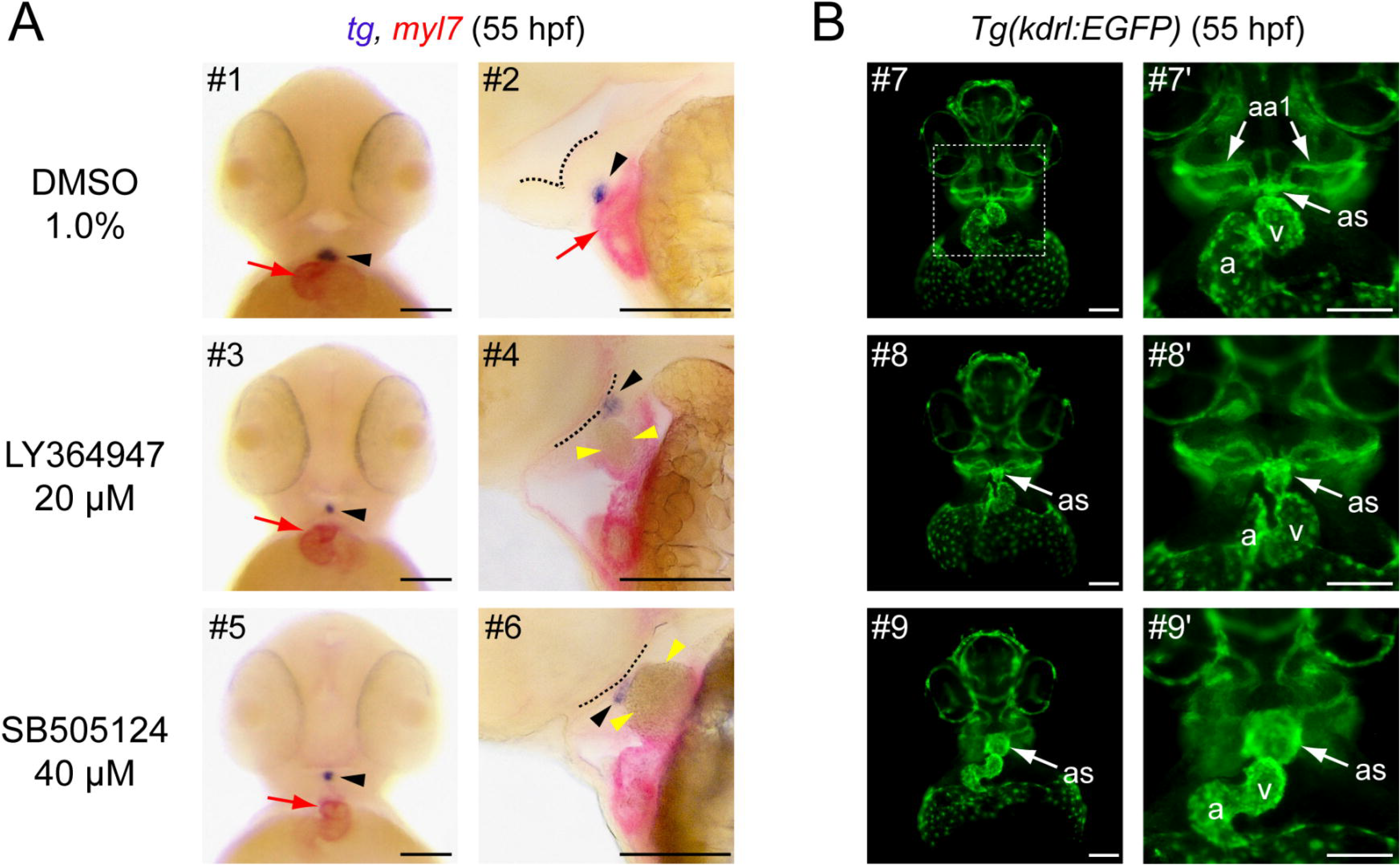
Ectopic thyroid positioning due to cardiac outflow tract malformation. **(A)** Dual-color *in situ* hybridization of thyroidal *tg* (black arrowheads) and cardiac *myl7* (red arrows) expression in 55 hpf embryos treated with Nodal/TGFβ inhibitors LY364947 and SB505124 from 24 to 54 hpf. The left panel of images show ventral views, anterior is to the top. The right panel of images shows mid-sagittal vibratome sections (anterior to the left) of stained embryos from the same experiment. Phenotypic comparison of inhibitor-treated and control embryos (DMSO, dimethylsulfoxide) revealed reduced thyroid size after treatment with both inhibitors (#3,5) as well as an apparent ectopic location of the thyroid primordium close to the pharyngeal floor (indicated as dotted line in #4,6). Note the apparent gap between thyroid marker expression (black arrowheads) and the distal portion of the *myl7*-expressing cardiac outflow tract in ventral views (see #3,5) and the large accumulation of blood cells (yellow arrowheads) in the distal part of the outflow tract as evident from vibratome sections (#4,6). **(B)** Transgenic *Tg(kdrl:EGFP)* embryos were treated from 24 to 54 hpf with Nodal/TGFβ inhibitors LY364947 and SB505124 and endothelial/endocardial GFP reporter expression was analyzed by immunofluorescence staining at 55 hpf. The left panel of epifluorescence images (#7,8,9) shows large field views of the head region (ventral view, anterior to the top) while the right panel shows magnified views (#7’,8’,9’) of the outflow tract region (corresponding to the boxed area shown for embryo #7). Note the massive ballooning of the aortic sac (as) in inhibitor-treated embryos as compared to control embryos. Abbreviations: a atrium; v ventricle; aa1 first pair of aortic arch arteries. Scale bars: 100 µm.

Pharyngula treatment with inhibitors of BMP (Fig. 8A), FGF (Fig. 8A) and TGFβ (Fig. 9A) signaling caused concentration-dependent decrements in thyroid size in 55 hpf embryos. Penetrance and severity of these effects was stronger for BMP and FGF inhibitors (Fig. 8C) with many treated embryos displaying only very tiny patches of *tg*-expressing cells (see embryos #4,6 in Fig. 8A). In the BMP inhibitor treatment groups, up to one third of embryos displayed supernumerary clusters of *tg*-expressing cells located bilaterally to the cardiac OFT (see #3 in Fig. 8A). Cardiac looping was not affected in these treatments but inhibition of BMP signaling was associated with an increased ventricle size compared to controls (see embryo #3 in Fig. 8A).

Treatment with inhibitors of FGF signaling caused a more complex phenotype. Treated embryos had a much smaller head size, a reduced head-trunk angle (35) indicative of delayed development and showed compromised morphogenesis of the pharyngeal region (Fig. 8B). Interestingly, treatment with the FGF inhibitor PD166866 resulted in delayed or even incomplete detachment of the thyroid primordium from the pharyngeal floor (Fig. 8B). From dual-color WISH analyses, we noted an increased space between the thyroid and the cardiac OFT indicating that inhibition of FGF signaling disrupted the normal coordination of thyroid budding and heart descent into the sub-pharyngeal region (Fig. 8B).

A similar budding defect appeared to occur following inhibition of Nodal/TGFβ signaling (Fig. 9A). During the initial phenotyping of double stained embryos, we observed a high proportion of Nodal/TGFβ inhibitor-treated embryos with a thyroid primordium positioned close to the pharyngeal floor but at large distance to *myl7*-expressing cardiac tissue (see embryos #3,5 in Fig. 9A). These embryos also presented mild pericardial edema. Subsequent vibratome sections of stained embryos revealed large accumulations of blood cells in the distal part of the OFT (see embryos #4,6 in Fig. 9A). When analyzing Nodal/TGFβ inhibitor-treated embryos carrying the *kdrl-EGFP* transgene to visualize endocardial and endothelial cells, we observed a massive ballooning of the aortic sac (Fig. 9B), a transient embryonic structure connecting the cardiac OFT to the first pair of aortic arch arteries and the ventral aorta (15). Since the aortic sac is not covered by a myocardial cell layer, this ballooning effect was not visualized by *myl7* staining. These observations indicated that the massively enlarged aortic sac was pushing the thyroid diverticulum towards the pharyngeal floor preventing an effective relocalization of the thyroid. Additional analyses of live *kdrl-EGFP* embryos indicated that the aortic sac ballooning and blood cell accumulation were due to defective blood flow (data not shown). Upon removal of the inhibitors from the medium at 55 hpf, we observed a gradual recovery of cardiac functionality, a concomitant regression of the aortic sac and, within 24 to 48 hours of drug withdrawal, relocalization of the thyroid into the hypobranchial region became evident (data not shown).

Although thyroids of cyclopamine-treated embryos at 55 hpf had a comparable size as their control counterparts, approximately half of the treated embryos displayed an aberrant thyroid morphology (Fig. 9). Most prevalent were uni- or bilateral expansions of the *tg* expression domain (see embryo #8 in Fig. 8A). Incidence of thyroid anomalies in cyclopamine-treated embryos increased concomitantly with the prevalence and severity of vascular alterations (data not shown). Co-occurrence of vascular and thyroidal abnormalities (irregularly enlarged thyroids) was observed also for embryos treated with the PDGF inhibitor KI11502 and VEGF inhibitors (vatalanib and SU4312). Disorganized and enlarged thyroids detected in these treatment groups resembled thyroid phenotypes described in embryos treated with these compounds in the somitogenesis screen (this study, see Fig. 5B) and in previous studies (15).

## Discussion

In this study, we employed a pharmacological strategy to gain insights into the role of major signaling pathways during early stages of zebrafish thyroid development. Given the experimentally supported concept that cardiac mesoderm provides a source of instructive signals for different steps of thyroid development (10, 11, 15, 26, 54, 55), our screening approach included a comprehensive phenotyping of drug-induced deviations of thyroid morphogenesis and concomitant cardiovascular anomalies. One advantage of pharmacological approaches is that they can afford a level of temporal control of pathway modulation that is often difficult to achieve by genetic model systems (32, 56). This is an important asset as the literature of endoderm organ development is rich in instances where specific signaling pathways have reiterative, diverging and even opposite roles during organ development depending on the temporal context (29, 57, 58). Analyses of thyroid phenotypes following timed small molecule treatment in the present study provide several such examples for the importance of developmental context. For instance, while FGF inhibition during gastrula stages caused enhanced thyroidal *nkx2.4b* expression at 28 hpf (see Fig. 2), application of the same FGF inhibitors during somitogenesis completely abolished expression of this early thyroid marker in the prospective thyroid region (see Fig. 6). For the purpose of this first screening study, we temporally restricted small molecule treatments to cover three important phases of early zebrafish development (gastrula, somitogenesis, pharyngula). We expect that results from these screens will provide valuable guidance for further temporal refinements of treatments periods in order to more precisely describe iterative pathway functions and identify critical developmental periods at a higher temporal resolution.

When probing the role of different pathways during gastrulation, thyroid phenotypes became evident following manipulation of four major signaling pathways (Wnt, FGF, BMP, Nodal/TGFβ) implicated in endoderm formation and early endodermal anterior-posterior patterning. For example, chemical blockage of FGF signaling, a pathway that normally exerts posteriorizing activity on endoderm patterning (59), caused enhanced thyroidal *nkx2.4b* expression in the anterior endoderm of zebrafish embryos. Such promoting effects on the specification of an anterior cell lineage fit well into concepts that drug-induced dampening of posteriorizing signals during gastrula promotes development of anterior tissue at the expense of posterior tissues (28). Corroborating this concept, we observed that treatment with FGF signaling inhibitors caused a dramatic truncation of the body at the level of the first somites (see Supplementary Fig. 2A), a gross phenotype reminiscent of what has previously been observed following dominant-negative FGF receptor overexpression (46, 60). However, still other morphogenetic effects might contributed to this specific thyroid phenotype given that early FGF blockage was previously shown to increase also total numbers of endodermal cells in *Xenopus* and zebrafish embryos (43, 61, 62).

Our gastrula screen faithfully recovered thyroid phenotypes for other pathways with reported posteriorizing effects on early anterior-posterior patterning of the endoderm (63). For example, over-activation of canonical Wnt signaling during gastrulation strongly suppressed TPC specification in the anterior endoderm. The contribution of posteriorizing activities of Wnt signaling to impaired TPC specification is further supported by a striking correlation between prevalence and severity of thyroid specification defects and neural posteriorization phenotypes in embryos treated with Wnt-activating compounds. Combined with the lack of prominent thyroid phenotypes following chemical blockage of early Wnt signaling, our observations indicate that endodermal progenitors of the thyroid cell lineage require an environment with low endogenous Wnt signaling during early development.

However, referring only to known anterior-posterior patterning activities of signaling pathways falls short in explaining some other phenotypes observed in our gastrula screen. For example, inhibition of BMP signaling in gastrula embryos severely impaired thyroid specification, a phenotype that is difficult to interpret in the light of reported posteriorizing activity of BMP on early endoderm patterning (64). Yet, it is important to note that BMP also plays a central role in dorsal-ventral (DV) patterning (65). Recent studies on early endoderm patterning showed that BMP is indeed required to promote ventral fates in the foregut endoderm while it represses dorsal fates (66). Although the zebrafish anterior endoderm at the time of thyroid specification is still a simple epithelial sheet lacking apparent morphological correlates of a DV axis (53), the thyroid bud will later develop from the ventral foregut endoderm. In the light of the reported down-regulation of ventral foregut markers in *Xenopus* embryos treated with the BMP inhibitor DMH1 (66), our findings in DMH1-treated zebrafish embryos suggest that a ventral identity might be assigned to the prospective endodermal thyroid progenitors already during gastrula stages and that such early DV patterning of the endoderm requires BMP.

Nodal/TGFβ signaling is yet another important, early acting pathway for endoderm formation and anterior-posterior axis patterning (67). During zebrafish blastula and early gastrula stages, Nodal activity is essential for the specification of mesendoderm in the marginal zone (68) and zebrafish mutants with compromised Nodal/TGFβ activity exhibit deficient endoderm formation along with severe thyroid specification defects (7, 31). However, when temporally restricting the inhibition of Nodal/TGFβ signaling to stages between 6 and 10 hpf, as in our gastrula screen, thyroid specification was only modestly affected. Notably, despite a marked reduction in the amount of anterior endoderm, inhibitor-treated embryos displayed only small reductions in the size of the thyroidal *nkx2.4b* expression domain. It appears, therefore, that the development of endodermal progenitors with a competence to differentiate into thyroid cells is largely independent of Nodal/TGFβ signaling in the period from 6 to 10 hpf. Conversely, the high prevalence of embryos displaying ectopic *nkx2.4b* expression at more posterior positions implies a possible posteriorizing activity of Nodal/TGFβ signaling during endodermal anterior-posterior patterning. Collectively, results from our gastrula screen indicate that Wnt and FGF signaling during gastrulation stages are dispensable for development of endodermal progenitors of the thyroid lineage whereas gastrula BMP signaling is absolutely required for later thyroid specification.

During somitogenesis, the pre-patterned endoderm undergoes further intricate patterning accompanied by the acquisition of regionalized cell fates. This process results in the definition of stereotypic regions at which endoderm-derived organ buds will be induced along the primitive gut tube. In contrast to the induction of hepatic, pancreatic and lung cells fates (24, 25, 29), very little is known about the identity of permissive and inductive signaling cues regulating thyroid cell specification. Thus, thyroid phenotypes arising from pathway manipulation during somitogenesis stages are of particular interest to fill this gap of knowledge. One key finding of our somitogenesis screen was a strict requirement of FGF and BMP signaling for thyroid cell specification to occur in zebrafish embryos. Inhibition of either pathway caused severe impairments of thyroid development and the thyroid phenotypes observed at high concentrations of BMP and FGF signaling inhibitors could be classified as true thyroid agenesis because of a complete lack of thyroid cell specification and later thyroid tissue formation. Moreover, it is of particular note that our somitogenesis screen identified with BMP and FGF the same two signaling pathways that are currently considered the most potent signals to induce thyroid cell differentiation in human and murine stem cell models (69, 70). This screening outcome underscores the value of the zebrafish model as a powerful *in vivo* system to interrogate signaling cues controling vertebrate thyroid organogenesis.

Several previous *in vivo* studies have documented a pivotal role of FGF signaling in regulating embryonic thyroid development in mouse (71–73), *Xenopus* (69, 74) and zebrafish (10). The thyroid cell specification defects observed in our somitogenesis screen following treatment with two different FGF inhibitors (PD166866, SU5402) are qualitatively similar to results from a previous zebrafish study using SU5402 treatment during somitogenesis (10). In zebrafish, fgf8 has been identified as one important factor for thyroid specification (10). Since zebrafish fgf8 is expressed in the cardiac mesoderm adjacent to the converging anterior endoderm throughout somitogenesis stages, this finding adds further evidence to the presumed critical role of cardiac mesoderm as a source of instructing factors for thyroid cell specification. Previous studies suggested that, although required for thyroid cell specification, FGF signaling might not exert inductive activity in zebrafish but rather acts as a permissive factor (10). This contention is supported by our observations that somitogenesis treatment with BCI, a compound that enhances local FGF signaling (75), did not cause expanded or ectopic *nkx2.4b* expression. Notably, although diverse defects during thyroid organogenesis were observed in murine genetic models with impaired FGF signaling (55, 72, 73), none of these genetic studies reported a severe thyroid specification defect as observed in zebrafish. It is likely that redundant activities among different branches of FGF signaling prevented such early thyroid phenotypes in genetic models. However, similar to what we observed in zebrafish embryos, global pharmacological FGF inhibition in mouse embryonic explant cultures effectively blocks induction of thyroidal Nkx2.1 expression in foregut endoderm (70).

In contrast to FGF, no data were previously available on the importance of BMP signaling for early thyroid development in zebrafish and no conclusive information has been obtained from genetic mouse models with perturbed BMP signaling. In *Twisted* null mice, for example, thyroidal expression of *Hhex* is reportedly absent at the 10-somite stage but these mice later develop macroscopically visible thyroid glands of unknown functionality (76). In mutant mice lacking the endogenous BMP inhibitor chordin, hypoplastic thyroids develop in a severely disorganized tracheal region but no data are available about the development of the early thyroid anlage in these mutants (77). While our somitogenesis screen results provide the first demonstration of an absolute requirement of BMP signaling for thyroid cell specification in zebrafish, these findings compare well to recent observations in *Xenopus* embryos (69) and mouse embryonic explant cultures (69, 70), corroborating an evolutionary conserved role of BMP for early thyroid development across vertebrates. Notably, in all three experimental systems, pharmacological inhibition of BMP signaling during somitogenesis stages completely blocked the induction of early thyroid marker expression in the prospective thyroid region.

Our screening revealed further that FGF and BMP are still required at later stages of thyroid morphogenesis. The presence of small and tiny thyroids in stage 55 hpf embryos treated with FGF inhibitors during the pharyngula period is reminiscent of the hypoplastic thyroid phenotypes reported for various murine models with impaired FGF signaling (55, 73). Moreover, similar to reports of anomalous thyroid budding in mouse embryos with defective FGF signaling (55), we confirmed that distorted thyroid budding is one component of the thyroid phenotype in FGF inhibitor-treated zebrafish embryos.

BMPs and FGFs are diffusible growth factors and several FGFs and BMPs are expressed in the cardiac mesoderm adjacent to the developing thyroid (55, 73, 78). While pharmacological inhibition of FGF and BMP signaling clearly results in failure of thyroid development, it is conceivable that perturbed development of cardiac mesoderm or abnormal positioning of cardiac mesoderm relative to the population of endodermal thyroid progenitors could similarly affect early thyroid development as a consequence of altered local signaling activities. Proof-of-principle experiments with embryonic foregut endoderm explants have shown that endoderm cultured in the absence of mesoderm fails to induce TPC specification whereas addition of exogenous FGF2 and BMP4 to mesoderm-depleted endoderm explants effectively induces early thyroid marker expression including *nkx2.1, pax2* and *hhex* (69).

In fact, our combined analyses of cardiac and thyroid development suggest that altered mesoderm-endoderm interactions might be involved in some thyroid phenotypes recovered in our screens. One such example is the thyroid agenesis caused by over-activation of canonical Wnt signaling during somitogenesis stages. While the overall normal thyroid anlage formation following IWR1 treatment indicates that Wnt signaling during somitogenesis is dispensable for thyroid specification, the complete lack of thyroid marker expression following treatment with Wnt inducers has, to our knowledge, no precedent in other model organisms. Wnt signaling is required for development of several endoderm-derived organs including pancreas, liver, and swim bladder in zebrafish (24, 58, 79) but the role of this pathway for thyroid induction has not yet been addressed.

At a first glance, thyroid specification phenotypes resulting from Wnt over-activation during gastrula and somitogenesis stages appear quite similar, but there are several notable differences. First, whereas gastrula stage-treated embryos displayed severe posteriorization phenotypes indicating thyroid loss as a consequence of early anterior-posterior patterning defects, such global posteriorization effects were less prominent in somitogenesis-treated embryos. Second, almost all embryos treated with Wnt inducers during gastrula stages displayed at least some small domains of *tg* expression at 55 hpf whereas Wnt over-activation during somitogenesis caused a permanent lack of thyroid marker expression. Third, combined analysis of cardiac and thyroid development revealed a very close correlation between diminished cardiac mesoderm formation and reduced thyroid marker expression. The latter observation is particularly intriguing as other zebrafish models with severely impaired cardiac formation also showed marked deficits in thyroid specification (10). Consistent with previous reports (57), we found that drug-induced over-activation of Wnt signaling at stages after heart tube formation no longer impairs thyroid marker expression. Further analyses of Wnt-induced phenotypes in cardiac and thyroid development are warranted to shed light on the underlying developmental mechanisms leading to thyroid agenesis.

Disturbed positioning of cardiac mesoderm relative to endodermal thyroid progenitors might also be involved in the expanded expression domain of the early thyroid marker *nkx2.4b* following inhibition of PDGF signaling. Although the PDGF receptor tyrosine kinase inhibitor KI11502 causing this thyroid specification effect is known to effectively phenocopy various developmental defects of *pdgfra* zebrafish mutants (50), a mechanistic interpretation of this phenotype remains quite puzzling. First, similar to the normal endoderm development reported in *pdgfra*-deficient zebrafish (50, 80), anterior endoderm formation was unaffected in KI11502-treated embryos. Second, while ligands of the PDGF receptor are expressed in anterior endoderm of zebrafish (*pdgfaa, pdgfab*) and in murine ventral foregut endoderm (*Pdgfa*), expression of PDGF receptors is essentially absent in the anterior endoderm during periods of thyroid specification in mouse and zebrafish (50, 81). Together, these findings point to a non-cell autonomous mechanism possibly involving interactions between endoderm and adjacent tissues. Accordingly, it was of particular interest that KI11502-treated embryos displayed a very specific cardiac developmental phenotype; a delay of midline fusion of the bilateral heart fields and heart tube assembly. Around the time of thyroid specification (23/24 hpf), this cardiac-specific delay resulted in a peculiar anatomical constellation where the prospective thyroid region within the anterior endoderm is juxtaposed to an enlarged cardiac field if compared to the anatomically more discrete and smaller arterial pole structures of the forming heart tube present at this stage in control embryos. One intriguing hypothesis is that due to the mispositioning of an enlarged field of cardiac mesoderm, an expanded region of anterior endoderm could be exposed to cardiac-borne signals with instructive capacity to induce endodermal *nkx2.4b* expression. Since KI11502 treatment effectively phenocopied the cardiac fusion defects of zebrafish *pdgfra* mutants (50, 80), such mutant models will provide an important resource for further experimental studies to verify or refute this hypothesis in genetic model systems.

During phenotyping of embryos at 55 hpf, we noticed that global manipulation of signaling pathways often results in grossly disorganized morphogenesis of the pharyngeal region including tissues surrounding the developing thyroid. This was particularly evident for the OFT region and the pharyngeal vasculature. Accordingly, caution is required in such cases when attempting to mechanistically link early signal perturbations to late stage thyroid phenotypes. Likely, certain late thyroid phenotypes could result from an overall maldevelopment of the tissue neighbo578rhood rather than from specific signaling perturbations acting directly on developing thyroid cells. In mouse and zebrafish, the nascent thyroid bud is associated with tissues of the cardiac OFT, most prominent is the direct apposition to the aortic sac (8, 15). A number of previous studies demonstrated that aberrant vessel development is often associated with irregular thyroid morphogenesis (11, 15, 54, 82) suggesting that the thyroid primordium is responsive to instructive cues from pharyngeal vessels influencing size, shape and localization of the diverticulum after detachment from the pharyngeal floor epithelium. Results from our screen reinforced this view as, in fact, most experimental treatments causing irregular and enlarged domains of *tg* expression in stage 55 hpf embryos showed concurrent defects in formation of the anterior arch arteries and the hypobranchial artery; i.e. vessels previously shown to guide positioning of thyroid tissue in zebrafish embryos and larvae (15, 82). The identification of a massively enlarged aortic sac due to cardiac dysfunction as the cause for a seemingly ectopic positioning of thyroid buds in TGFβ inhibitor-treated embryos further underscored the value of a combined thyroid and cardiovascular phenotyping approach for enhanced phenotype interpretation.

We also identified signaling pathways that appeared to have, if any, only a subordinate role in early zebrafish thyroid development. Consistent with conclusions reached in previous mouse and *Xenopus* studies (11, 17, 54, 83), results from our three screens indicate that Shh signaling is dispensable for thyroid specification. If detected, abnormalities in thyroid anlage formation coincided with impaired anterior endoderm formation (somitogenesis screen) and treatment with potent inducers of Shh signaling (i.e. purmorphamine) did not evoke detectable thyroid abnormalities in any of our screens. Moreover, cyclopamine-induced thyroid abnormalities present in later stage embryos always occurred along with defective cardiac OFT and pharyngeal vessel morphogenesis.

Thyroid phenotyping of embryos in all three screens failed to detect effects of Notch pathway inhibition on early zebrafish thyroid development indicating if any only a minor role of Notch-mediated lateral inhibition in the regulation of thyroid anlage specification and thyroid primordium formation during normal development. Consistent results were obtained using two new generation Notch pathway inhibitors, LY411575 and RO4929097, both of which inhibit γ-secretase activity, an accepted pharmacological approach to effectively block the Notch pathway in multiple model systems (84). Embryos treated with LY411575 and RO4929097 in our screens displayed well documented phenotypes of Notch pathway inhibition (85, 86) indicating that Notch signaling was effectively blocked by our experimental conditions. One previous zebrafish study reported significant increases in the size of the thyroid anlage (24 hpf) and the thyroid primordium (48 hpf) in Delta-Notch signaling-deficient *mindbomb* mutants compared to their WT siblings and significantly enlarged thyroid primordia following treatment with another γ-secretase inhibitor DAPT (87). Although the reasons for the conflicting findings are currently unknown, it remains possible that the prominent pharyngeal vessel anomalies detected in *mindbomb* mutants and DAPT-treated embryos contributed to a stronger thyroid phenotype. In our experiments, only few embryos treated with high concentrations of RO4929097 showed mild thyroid anomalies, cardiac edema and vascular defects at stage 55 hpf. Given recent data on a possible association of defective *JAG1*-Notch signaling with congenital hypothyroidism (88), further experimental studies are needed to examine NOTCH-dependent developmental processes during thyroid morphogenesis.

In summary, our studies showed that small molecule screening approaches are a valuable tool to uncover the role of major signaling pathways during thyroid morphogenesis in zebrafish. Our pharmacological screens identified BMP and FGF signaling as key factors for thyroid specification and early thyroid organogenesis in zebrafish, revealed the importance of low Wnt activities during early development for normal thyroid specification and corroborated further evidence for the contention that pharyngeal vessels provide guidance for size, shape and localization of thyroid tissue. Moreover, we identified several cases where the specific thyroid phenotype is likely due to perturbed local signaling as a result of defective cardiac mesoderm development. Given the plentiful signaling biosensor models available to the zebrafish community (89), studies are now required to map sites of action for critical pathways such as BMP and FGF signaling during early zebrafish development. By comparing our results with available data from other model organisms including *Xenopus*, chicken and mouse, we found that signaling cues regulating thyroid development appear broadly conserved across vertebrates. Thus, we expect that findings in the zebrafish model can inform mammalian models of thyroid organogenesis to further our understanding of the molecular mechanisms of congenital thyroid diseases. Moreover, we are confident that our data sets will provide a valuable resource enabling the formulation of new and testable hypotheses with respect to patterning cues that could improve the efficiency of *in vitro* stem cell-based protocols to generate thyroid cells.

## Acknowledgments

We thank V. Janssens for zebrafish husbandry and J.-M. Vanderwinden from the Light Microscopy Facility for technical assistance. This work was supported by grants from the Belgian National Fund for Scientific Research (FNRS) (FRSM 3-4598-12; CDR-J.0145.16), the Action de Recherche Concertée (ARC) de la Communauté Française de Belgique (ARC AUWB-2012-12/17-ULB3), the Fonds d’Encouragement à la Recherche de l’Université Libre de Bruxelles (FER-ULB), the Fund Yvonne Smits (King Baudouin Fundation) and the Berlin Institute of Health (BIH, CRG-TP2). B.H., P.G, and N.G are Fund for Research in the Industry and the Agriculture (FRIA) research fellows; S.C. is an FNRS Senior Research Associate.

## Additional Information

Supplementary information accompanies this paper.

## References

1. Holzer G, Laudet V 2013 Thyroid hormones and postembryonic development in amniotes. Curr Top Dev Biol 103:397–425.

2. De Felice M, Di Lauro R 2004 Thyroid development and its disorders: genetics and molecular mechanisms. Endocr Rev 25:722–746.

3. Nilsson M, Fagman H 2017 Development of the thyroid gland. Development 144:2123– 2140.

4. Fagman H, Nilsson M 2010 Morphogenesis of the thyroid gland. Mol Cell Endocrinol 323:35–54.

5. Fernandez LP, Lopez-Marquez A, Santisteban P 2015 Thyroid transcription factors in development, differentiation and disease. Nat Rev Endocrinol 11:29–42.

6. De Felice M, Di Lauro R 2011 Minireview: Intrinsic and extrinsic factors in thyroid gland development: an update. Endocrinology 152:2948–2956.

7. Elsalini OA, von Gartzen J, Cramer M, Rohr KB 2003 Zebrafish hhex, nk2.1a, and pax2.1 regulate thyroid growth and differentiation downstream of Nodal-dependent transcription factors. Dev Biol 263:67–80.

8. Fagman H, Andersson L, Nilsson M 2006 The developing mouse thyroid: embryonic vessel contacts and parenchymal growth pattern during specification, budding, migration, and lobulation. Dev Dyn 235:444–455.

9. Trueba SS, Auge J, Mattei G, Etchevers H, Martinovic J, Czernichow P, Vekemans M, Polak M, Attie-Bitach T 2005 PAX8, TITF1, and FOXE1 gene expression patterns during human development: new insights into human thyroid development and thyroid dysgenesis-associated malformations. J Clin Endocrinol Metab 90:455–462.

10. Wendl T, Adzic D, Schoenebeck JJ, Scholpp S, Brand M, Yelon D, Rohr KB 2007 Early developmental specification of the thyroid gland depends on han-expressing surrounding tissue and on FGF signals. Development 134:2871–2879.

11. Alt B, Elsalini OA, Schrumpf P, Haufs N, Lawson ND, Schwabe GC, Mundlos S, Gruters A, Krude H, Rohr KB 2006 Arteries define the position of the thyroid gland during its developmental relocalisation. Development 133:3797–3804.

12. Szinnai G, Lacroix L, Carre A, Guimiot F, Talbot M, Martinovic J, Delezoide A-L, Vekemans M, Michiels S, Caillou B, Schlumberger M, Bidart J-M, Polak M 2007 Sodium/iodide symporter (NIS) gene expression is the limiting step for the onset of thyroid function in the human fetus. J Clin Endocrinol Metab 92:70–76.

13. Alt B, Reibe S, Feitosa NM, Elsalini OA, Wendl T, Rohr KB 2006 Analysis of origin and growth of the thyroid gland in zebrafish. Dev Dyn 235:1872–1883.

14. Opitz R, Maquet E, Zoenen M, Dadhich R, Costagliola S 2011 TSH receptor function is required for normal thyroid differentiation in zebrafish. Mol Endocrinol 25:1579–1599.

15. Opitz R, Maquet E, Huisken J, Antonica F, Trubiroha A, Pottier G, Janssens V, Costagliola S 2012 Transgenic zebrafish illuminate the dynamics of thyroid morphogenesis and its relationship to cardiovascular development. Dev Biol 372:203–216.

16. Maeda K, Asai R, Maruyama K, Kurihara Y, Nakanishi T, Kurihara H, Miyagawa-Tomita S 2016 Postotic and preotic cranial neural crest cells differently contribute to thyroid development. Dev Biol 409:72–83.

17. Parlato R, Rosica A, Rodriguez-Mallon A, Affuso A, Postiglione MP, Arra C, Mansouri A, Kimura S, Di Lauro R, De Felice M 2004 An integrated regulatory network controlling survival and migration in thyroid organogenesis. Dev Biol 276:464–475.

18. Trubiroha A, Gillotay P, Giusti N, Gacquer D, Libert F, Lefort A, Haerlingen B, De Deken X, Opitz R, Costagliola S 2018 A Rapid CRISPR/Cas-based Mutagenesis Assay in Zebrafish for Identification of Genes Involved in Thyroid Morphogenesis and Function. Sci Rep 8:5647.

19. Fagman H, Nilsson M 2011 Morphogenetics of early thyroid development. J Mol Endocrinol 46:R33–42.

20. Fagman H, Liao J, Westerlund J, Andersson L, Morrow BE, Nilsson M 2007 The 22q11 deletion syndrome candidate gene Tbx1 determines thyroid size and positioning. Hum Mol Genet 16:276–285.

21. Nilsson M, Fagman H 2013 Mechanisms of thyroid development and dysgenesis: an analysis based on developmental stages and concurrent embryonic anatomy. Curr Top Dev Biol 106:123–170.

22. Porazzi P, Calebiro D, Benato F, Tiso N, Persani L 2009 Thyroid gland development and function in the zebrafish model. Mol Cell Endocrinol 312:14–23.

23. Lammert E, Cleaver O, Melton D 2001 Induction of pancreatic differentiation by signals from blood vessels. Science 294:564–567.

24. Ober EA, Verkade H, Field HA, Stainier DYR 2006 Mesodermal Wnt2b signalling positively regulates liver specification. Nature 442:688–691.

25. Manfroid I, Delporte F, Baudhuin A, Motte P, Neumann CJ, Voz ML, Martial JA, Peers B 2007 Reciprocal endoderm-mesoderm interactions mediated by fgf24 and fgf10 govern pancreas development. Development 134:4011–4021.

26. Serls AE, Doherty S, Parvatiyar P, Wells JM, Deutsch GH 2005 Different thresholds of fibroblast growth factors pattern the ventral foregut into liver and lung. Development 132:35–47.

27. Angelo JR, Tremblay KD 2018 Identification and fate mapping of the pancreatic mesenchyme. Dev Biol 435:15–25.

28. McCracken KW, Wells JM 2012 Molecular pathways controlling pancreas induction. Semin Cell Dev Biol 23:656–662.

29. Rankin SA, McCracken KW, Luedeke DM, Han L, Wells JM, Shannon JM, Zorn AM 2018 Timing is everything: Reiterative Wnt, BMP and RA signaling regulate developmental competence during endoderm organogenesis. Dev Biol 434:121–132.

30. Opitz R, Antonica F, Costagliola S 2013 New Model Systems to Illuminate Thyroid Organogenesis. Part I: An Update on the Zebrafish Toolbox. Eur Thyroid J 2:229–242.

31. Wendl T, Lun K, Mione M, Favor J, Brand M, Wilson SW, Rohr KB 2002 Pax2.1 is required for the development of thyroid follicles in zebrafish. Development 129:3751– 3760.

32. Kaufman CK, White RM, Zon L 2009 Chemical genetic screening in the zebrafish embryo. Nat Protoc 4:1422–1432.

33. Tamplin OJ, White RM, Jing L, Kaufman CK, Lacadie SA, Li P, Taylor AM, Zon LI 2012 Small molecule screening in zebrafish: swimming in potential drug therapies. Wiley Interdiscip Rev Dev Biol 1:459–468.

34. Westerfield M 2000 The zebrafish book. A guide for the laboratory use of zebrafish (Danio rerio). Univ. of Oregon Press.

35. Kimmel CB, Ballard WW, Kimmel SR, Ullmann B, Schilling TF 1995 Stages of embryonic development of the zebrafish. Dev Dyn 203:253–310.

36. White RM, Sessa A, Burke C, Bowman T, LeBlanc J, Ceol C, Bourque C, Dovey M, Goessling W, Burns CE, Zon LI 2008 Transparent adult zebrafish as a tool for in vivo transplantation analysis. Cell Stem Cell 2:183–189.

37. Mizoguchi T, Verkade H, Heath JK, Kuroiwa A, Kikuchi Y 2008 Sdf1/Cxcr4 signaling controls the dorsal migration of endodermal cells during zebrafish gastrulation. Development 135:2521–2529.

38. Huang C-J, Tu C-T, Hsiao C-D, Hsieh F-J, Tsai H-J 2003 Germ-line transmission of a myocardium-specific GFP transgene reveals critical regulatory elements in the cardiac myosin light chain 2 promoter of zebrafish. Dev Dyn 228:30–40.

39. Jin S-W, Beis D, Mitchell T, Chen J-N, Stainier DYR 2005 Cellular and molecular analyses of vascular tube and lumen formation in zebrafish. Development 132:5199–5209.

40. Yelon D, Horne SA, Stainier DY 1999 Restricted expression of cardiac myosin genes reveals regulated aspects of heart tube assembly in zebrafish. Dev Biol 214:23–37.

41. Thisse C, Thisse B 2008 High-resolution in situ hybridization to whole-mount zebrafish embryos. Nat Protoc 3:59–69.

42. Jowett T 2001 Double in situ hybridization techniques in zebrafish. Methods 23:345– 358.

43. Poulain M, Furthauer M, Thisse B, Thisse C, Lepage T 2006 Zebrafish endoderm formation is regulated by combinatorial Nodal, FGF and BMP signalling. Development 133:2189–2200.

44. Rohde LA, Heisenberg C-P 2007 Zebrafish gastrulation: cell movements, signals, and mechanisms. Int Rev Cytol 261:159–192.

45. Sanvitale CE, Kerr G, Chaikuad A, Ramel M-C, Mohedas AH, Reichert S, Wang Y, Triffitt JT, Cuny GD, Yu PB, Hill CS, Bullock AN 2013 A new class of small molecule inhibitor of BMP signaling. PloS One 8:e62721.

46. Griffin K, Patient R, Holder N 1995 Analysis of FGF function in normal and no tail zebrafish embryos reveals separate mechanisms for formation of the trunk and the tail. Development 121:2983–2994.

47. Sampath K, Rubinstein AL, Cheng AM, Liang JO, Fekany K, Solnica-Krezel L, Korzh V, Halpern ME, Wright CV 1998 Induction of the zebrafish ventral brain and floorplate requires cyclops/nodal signalling. Nature 395:185–189.

48. van de Water S, van de Wetering M, Joore J, Esseling J, Bink R, Clevers H, Zivkovic D 2001 Ectopic Wnt signal determines the eyeless phenotype of zebrafish masterblind mutant. Development 128:3877–3888.

49. Ye D, Xie H, Hu B, Lin F 2015 Endoderm convergence controls subduction of the myocardial precursors during heart-tube formation. Development 142:2928–2940.

50. Bloomekatz J, Singh R, Prall OW, Dunn AC, Vaughan M, Loo C-S, Harvey RP, Yelon D 2017 Platelet-derived growth factor (PDGF) signaling directs cardiomyocyte movement toward the midline during heart tube assembly. eLife 6.

51. Brand M, Heisenberg CP, Warga RM, Pelegri F, Karlstrom RO, Beuchle D, Picker A, Jiang YJ, Furutani-Seiki M, van Eeden FJ, Granato M, Haffter P, Hammerschmidt M, Kane DA, Kelsh RN, Mullins MC, Odenthal J, Nusslein-Volhard C 1996 Mutations affecting development of the midline and general body shape during zebrafish embryogenesis. Development 123:129–142.

52. Hami D, Grimes AC, Tsai H-J, Kirby ML 2011 Zebrafish cardiac development requires a conserved secondary heart field. Development 138:2389–2398.

53. Wallace KN, Pack M 2003 Unique and conserved aspects of gut development in zebrafish. Dev Biol 255:12–29.

54. Fagman H, Grande M, Gritli-Linde A, Nilsson M 2004 Genetic deletion of sonic hedgehog causes hemiagenesis and ectopic development of the thyroid in mouse. Am J Pathol 164:1865–1872.

55. Lania G, Zhang Z, Huynh T, Caprio C, Moon AM, Vitelli F, Baldini A 2009 Early thyroid development requires a Tbx1-Fgf8 pathway. Dev Biol 328:109–117.

56. Wiley DS, Redfield SE, Zon LI 2017 Chemical screening in zebrafish for novel biological and therapeutic discovery. Methods Cell Biol 138:651–679.

57. Shin D, Lee Y, Poss KD, Stainier DYR 2011 Restriction of hepatic competence by Fgf signaling. Development 138:1339–1348.

58. Goessling W, North TE, Lord AM, Ceol C, Lee S, Weidinger G, Bourque C, Strijbosch R, Haramis A-P, Puder M, Clevers H, Moon RT, Zon LI 2008 APC mutant zebrafish uncover a changing temporal requirement for wnt signaling in liver development. Dev Biol 320:161–174.

59. Dessimoz J, Opoka R, Kordich JJ, Grapin-Botton A, Wells JM 2006 FGF signaling is necessary for establishing gut tube domains along the anterior-posterior axis in vivo. Mech Dev 123:42–55.

60. Griffin KJP, Kimelman D 2003 Interplay between FGF, one-eyed pinhead, and T-box transcription factors during zebrafish posterior development. Dev Biol 264:456–466.

61. Cha S-W, Hwang Y-S, Chae J-P, Lee S-Y, Lee H-S, Daar I, Park MJ, Kim J 2004 Inhibition of FGF signaling causes expansion of the endoderm in Xenopus. Biochem Biophys Res Commun 315:100–106.

62. Mizoguchi T, Izawa T, Kuroiwa A, Kikuchi Y 2006 Fgf signaling negatively regulates Nodal-dependent endoderm induction in zebrafish. Dev Biol 300:612–622.

63. McLin VA, Rankin SA, Zorn AM 2007 Repression of Wnt/beta-catenin signaling in the anterior endoderm is essential for liver and pancreas development. Development 134:2207–2217.

64. Tiso N, Filippi A, Pauls S, Bortolussi M, Argenton F 2002 BMP signalling regulates anteroposterior endoderm patterning in zebrafish. Mech Dev 118:29–37.

65. Tucker JA, Mintzer KA, Mullins MC 2008 The BMP signaling gradient patterns dorsoventral tissues in a temporally progressive manner along the anteroposterior axis. Dev Cell 14:108–119.

66. Stevens ML, Chaturvedi P, Rankin SA, Macdonald M, Jagannathan S, Yukawa M, Barski A, Zorn AM 2017 Genomic integration of Wnt/beta-catenin and BMP/Smad1 signaling coordinates foregut and hindgut transcriptional programs. Development 144:1283–1295.

67. Tian T, Meng AM 2006 Nodal signals pattern vertebrate embryos. Cell Mol Life Sci 63:672–685.

68. Hagos EG, Dougan ST 2007 Time-dependent patterning of the mesoderm and endoderm by Nodal signals in zebrafish. BMC Dev Biol 7:22.

69. Kurmann AA, Serra M, Hawkins F, Rankin SA, Mori M, Astapova I, Ullas S, Lin S, Bilodeau M, Rossant J, Jean JC, Ikonomou L, Deterding RR, Shannon JM, Zorn AM, Hollenberg AN, Kotton DN 2015 Regeneration of Thyroid Function by Transplantation of Differentiated Pluripotent Stem Cells. Cell Stem Cell 17:527–542.

70. Serra M, Alysandratos K-D, Hawkins F, McCauley KB, Jacob A, Choi J, Caballero IS, Vedaie M, Kurmann AA, Ikonomou L, Hollenberg AN, Shannon JM, Kotton DN 2017 Pluripotent stem cell differentiation reveals distinct developmental pathways regulating lungversus thyroid-lineage specification. Development 144:3879–3893.

71. Celli G, LaRochelle WJ, Mackem S, Sharp R, Merlino G 1998 Soluble dominant-negative receptor uncovers essential roles for fibroblast growth factors in multi-organ induction and patterning. EMBO J 17:1642–1655.

72. Kameda Y, Ito M, Nishimaki T, Gotoh N 2009 FRS2alpha is required for the separation, migration, and survival of pharyngeal-endoderm derived organs including thyroid, ultimobranchial body, parathyroid, and thymus. Dev Dyn 238:503–513.

73. Liang S, Johansson E, Barila G, Altschuler DL, Fagman H, Nilsson M 2018 A branching morphogenesis program governs embryonic growth of the thyroid gland. Development 145.

74. Shifley ET, Kenny AP, Rankin SA, Zorn AM 2012 Prolonged FGF signaling is necessary for lung and liver induction in Xenopus. BMC Dev Biol 12:27.

75. Molina G, Vogt A, Bakan A, Dai W, Queiroz de Oliveira P, Znosko W, Smithgall TE, Bahar I, Lazo JS, Day BW, Tsang M 2009 Zebrafish chemical screening reveals an inhibitor of Dusp6 that expands cardiac cell lineages. Nat Chem Biol 5:680–687.

76. Petryk A, Anderson RM, Jarcho MP, Leaf I, Carlson CS, Klingensmith J, Shawlot W, O’Connor MB 2004 The mammalian twisted gastrulation gene functions in foregut and craniofacial development. Dev Biol 267:374–386.

77. Bachiller D, Klingensmith J, Shneyder N, Tran U, Anderson R, Rossant J, De Robertis EM 2003 The role of chordin/Bmp signals in mammalian pharyngeal development and DiGeorge syndrome. Development 130:3567–3578.

78. Danesh SM, Villasenor A, Chong D, Soukup C, Cleaver O 2009 BMP and BMP receptor expression during murine organogenesis. Gene Expr Patterns GEP 9:255–265.

79. Poulain M, Ober EA 2011 Interplay between Wnt2 and Wnt2bb controls multiple steps of early foregut-derived organ development. Development 138:3557–3568.

80. El-Rass S, Eisa-Beygi S, Khong E, Brand-Arzamendi K, Mauro A, Zhang H, Clark KJ, Ekker SC, Wen X-Y 2017 Disruption of pdgfra alters endocardial and myocardial fusion during zebrafish cardiac assembly. Biol Open 6:348–357.

81. Orr-Urtreger A, Lonai P 1992 Platelet-derived growth factor-A and its receptor are expressed in separate, but adjacent cell layers of the mouse embryo. Development 115:1045–1058.

82. Opitz R, Hitz M-P, Vandernoot I, Trubiroha A, Abu-Khudir R, Samuels M, Desilets V, Costagliola S, Andelfinger G, Deladoey J 2015 Functional zebrafish studies based on human genotyping point to netrin-1 as a link between aberrant cardiovascular development and thyroid dysgenesis. Endocrinology 156:377–388.

83. Rankin SA, Han L, McCracken KW, Kenny AP, Anglin CT, Grigg EA, Crawford CM, Wells JM, Shannon JM, Zorn AM 2016 A Retinoic Acid-Hedgehog Cascade Coordinates Mesoderm-Inducing Signals and Endoderm Competence during Lung Specification. Cell Rep 16:66–78.

84. Olsauskas-Kuprys R, Zlobin A, Osipo C 2013 Gamma secretase inhibitors of Notch signaling. OncoTargets Ther 6:943–955.

85. Arslanova D, Yang T, Xu X, Wong ST, Augelli-Szafran CE, Xia W 2010 Phenotypic analysis of images of zebrafish treated with Alzheimer’s gamma-secretase inhibitors. BMC Biotechnol 10:24.

86. Wang Y, Pan L, Moens CB, Appel B 2014 Notch3 establishes brain vascular integrity by regulating pericyte number. Development 141:307–317.

87. Porazzi P, Marelli F, Benato F, de Filippis T, Calebiro D, Argenton F, Tiso N, Persani L 2012 Disruptions of global and JAGGED1-mediated notch signaling affect thyroid morphogenesis in the zebrafish. Endocrinology 153:5645–5658.

88. de Filippis T, Marelli F, Nebbia G, Porazzi P, Corbetta S, Fugazzola L, Gastaldi R, Vigone MC, Biffanti R, Frizziero D, Mandara L, Prontera P, Salerno M, Maghnie M, Tiso N, Radetti G, Weber G, Persani L 2016 JAG1 Loss-Of-Function Variations as a Novel Predisposing Event in the Pathogenesis of Congenital Thyroid Defects. J Clin Endocrinol Metab 101:861–870.

89. Moro E, Vettori A, Porazzi P, Schiavone M, Rampazzo E, Casari A, Ek O, Facchinello N, Astone M, Zancan I, Milanetto M, Tiso N, Argenton F 2013 Generation and application of signaling pathway reporter lines in zebrafish. Mol Genet Genomics 288:231–242.

