## Supplementary Information for "Small Molecule Screening in Zebrafish Embryos Identifies Signaling Pathways Regulating Early Thyroid Development"

**This file contains 1 Supplementary Table and 5 Supplementary Figures**

**Supplementary Table 1**. List of small molecule compounds used in this study

**Supplementary Figure 1**. Induction of additional ectopic domains of *nkx2.4b* expression following inhibition of FGF signaling during zebrafish gastrulation.

**Supplementary Figure 2**. Aberrant morphology of *tg* expression domains following inhibition of FGF and Wnt signaling during zebrafish gastrulation.

**Supplementary Figure 3**. Induction of additional ectopic domains of *nkx2.4b* expression following inhibition of Nodal/TGFβ signaling during zebrafish gastrulation.

**Supplementary Figure 4**. Inhibition of PDGF signaling during somitogenesis delays cardiac fusion and causes abnormal heart tube assembly.

**Supplementary Figure 5**. Inhibition of Shh signaling during somitogenesis affects anterior endoderm formation.

### Supplementary Tables

**Supplementary Table 1.** List of small molecule compounds used in this study.

| Pathway | Compound | Supplier | Catalog # |
|---|---|---|---|
| BMP | DMH1 | Sigma | D8946 |
| BMP | LDN193189 | Selleck Chemicals | S2618 |
| FGF | SU5402 | Tocris | 3300 |
| FGF | PD166866 | Sigma | PZ0114 |
| FGF | (E/Z)-BCI hydrochloride | Sigma | B4313 |
| Wnt | IWR1 | Sigma | I0161 |
| Wnt | azakenpaullone | Sigma | A3734 |
| Wnt | BIO | Sigma | B1686 |
| Nodal/TGFβ | SB505124 | Sigma | S4696 |
| Nodal/TGFβ | LY364947 | Tocris | 2718 |
| Shh | cyclopamine | Merck-Millipore | 239803 |
| Shh | purmorphamine | Merck-Millipore | 540220 |
| Shh | SAG | Merck-Millipore | 566660 |
| PDGF | KI11502 | Merck-Millipore | 521234 |
| PDGF | AG1295 | Merck-Millipore | 658550 |
| VEGF | vatalanib | LCC | V-8303 |
| VEGF | SU4312 | Sigma | S8567 |
| VEGF | GS4012 | Merck-Millipore | 676491 |
| Notch | LY411575 | Sigma | SML0506 |
| Notch | RO4929097 | Selleck Chemicals | S1575 |

### Supplementary Figures 1 – 5

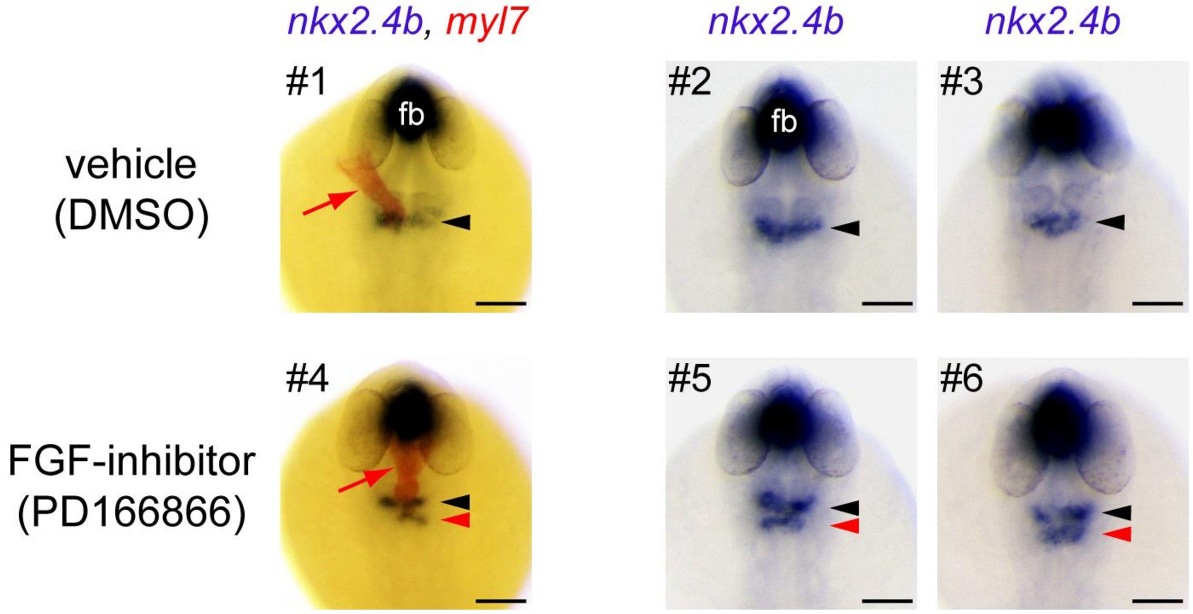


**Supplementary Figure 1. Induction of additional ectopic domains of *nkx2.4b* expression following inhibition of FGF signaling during zebrafish gastrulation.** Embryos treated from 6 to 10 hpf (gastrula screen) with small molecule inhibitors of FGF signaling were analyzed at 28 hpf for expression of the early thyroid marker *nkx2.4b* by *in situ* hybridization. Note that *nkx2.4b* is expressed in the thyroid anlage (arrowheads) and the ventral forebrain (fb). Dual-color staining with a *myl7* riboprobe (see #1,4) highlights the primitive heart tube (red arrows). Compared to embryos from the vehicle control group (0.5% DMSO), treatments with the FGF inhibitor PD166866 (8 µM) resulted in the induction of additional ectopic domains of *nkx2.4b* expression (red arrowheads) posterior to the orthotopic position of the thyroid anlage (black arrowheads). Dorsal views are shown, anterior is to the top. Scale bars: 100 µm.

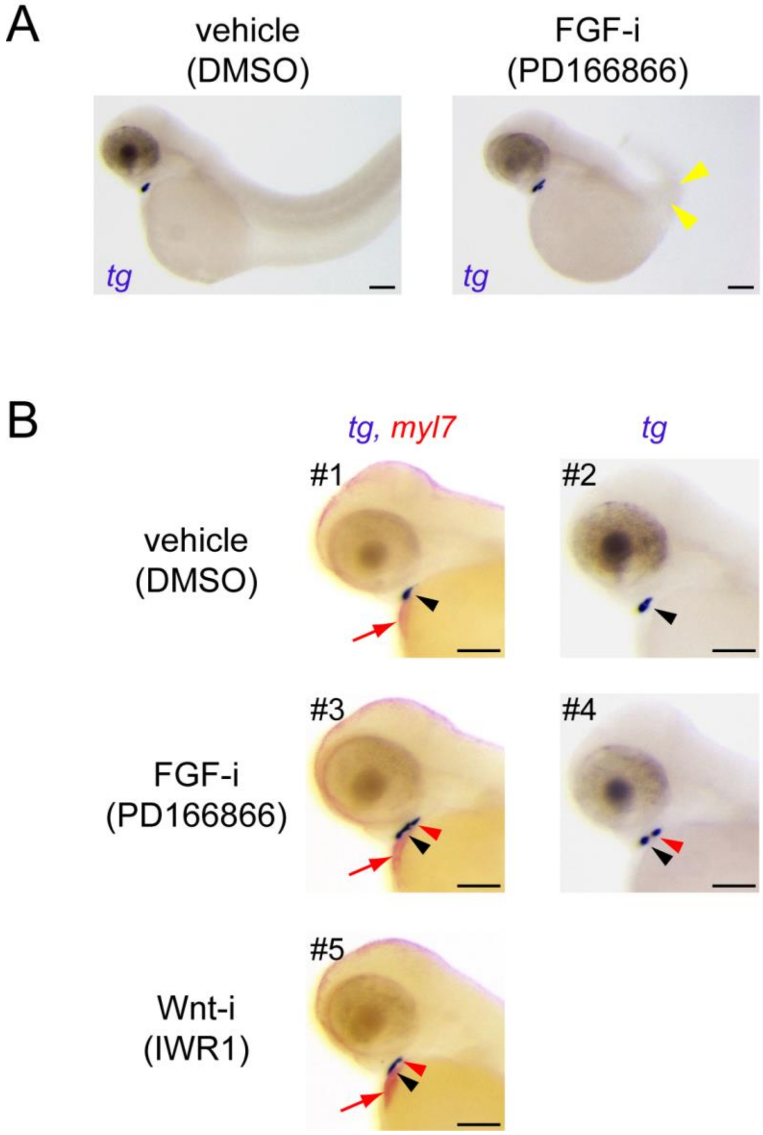


**Supplementary Figure 2. Aberrant morphology of *tg* expression domains following inhibition of FGF and Wnt signaling during zebrafish gastrulation.** Embryos treated from 6 to 10 hpf (gastrula screen) with small molecule inhibitors (i) of FGF and Wnt signaling were analyzed at 55 hpf for *tg* expression (arrowheads) by *in situ* hybridization. Dual-color staining with a *myl7* riboprobe (see #3,5) highlights the myocardium (red arrows). **(A)** Large field lateral views (anterior to the left) show the dramatic loss of caudal tissue (yellow arrowheads) in embryos treated with the FGF inhibitor PD166866 (8 µM). Scale bars: 200 µm. **(B)** Lateral views of the head region (anterior to the left) of 55 hpf embryos show aberrant morphologies of the *tg* expression domain in embryos treated with the FGF inhibitor PD166866 (8 µM) and the inhibitor of Wnt signaling IWR1 (20 µM). While control thyroids typically assume a drop-like shape (see #1,2), thyroids from inhibitor-treated embryos displayed an elongated shape with an expansion of the *tg* expression domain towards more posterior positions (red arrowheads). Scale bars: 100 µm.

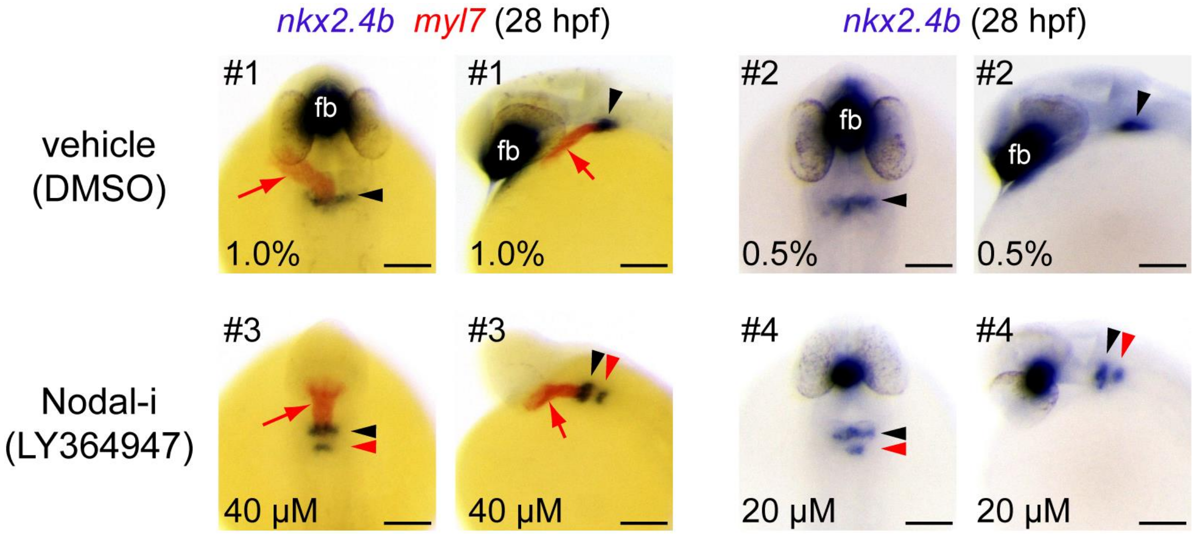


**Supplementary Figure 3. Induction of additional ectopic domains of *nkx2.4b* expression following inhibition of Nodal/TGFβ signaling during zebrafish gastrulation.** Embryos treated from 6 to 10 hpf (gastrula screen) with small molecule inhibitors (i) of Nodal/TGFβ signaling were analyzed at 28 hpf for expression of the early thyroid marker *nkx2.4b* by *in situ* hybridization. Note that *nkx2.4b* is expressed in the thyroid anlage (arrowheads) and the ventral forebrain (fb). Dual-color staining with a *myl7* riboprobe (see #1,3) highlights the primitive heart tube (red arrows). For each embryo, dorsal and lateral views are shown. Anterior is to the top and the left, respectively. Compared to embryos from the vehicle control group (1% DMSO), treatment with the Nodal/TGFβ inhibitor LY364947 resulted in the induction of additional ectopic domains of *nkx2.4b* expression (red arrowheads) posterior to the orthotopic position of the thyroid anlage (black arrowheads). Scale bars: 100 µm.

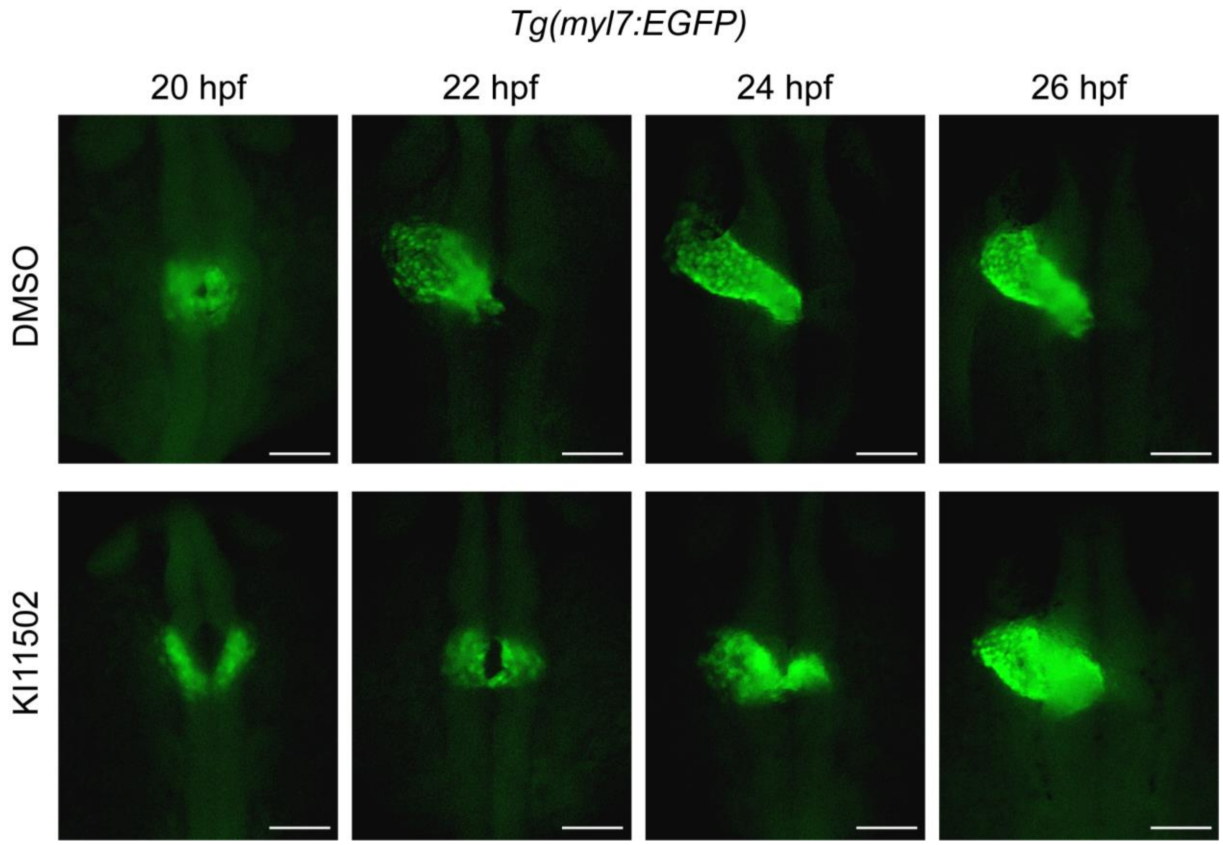


**Supplementary Figure 4. Inhibition of PDGF signaling during somitogenesis delays cardiac fusion and causes abnormal heart tube assembly.** Transgenic *Tg(myl7:EGFP)* embryos were treated from 10 up to 26 hpf with KI11502, an inhibitor of PDGF signaling, and GFP reporter expression was analyzed by immunofluorescence staining between 20 and 26 hpf to assess the time course of cardiac cone formation and assembly of the primitive heart tube. Examination of stage-matched 20 hpf embryos from control and inhibitor treatment groups showed a specific delay in the midline fusion of the converging bilateral heart fields in many embryos treated with 2 µM of KI11502. Although cardiac fusion occurred in KI11502-treated embryos in the following hours, heart tube assembly was compromised in many embryos (see for example the bifurcating heart fields at 24 hpf). By 26 hpf, KI11502-treated embryos displayed a rudimentary heart tube of greatly reduced anterior-posterior length and a broadened apical pole region. Dorsal views are shown, anterior is to the top. Scale bars: 100 µm.

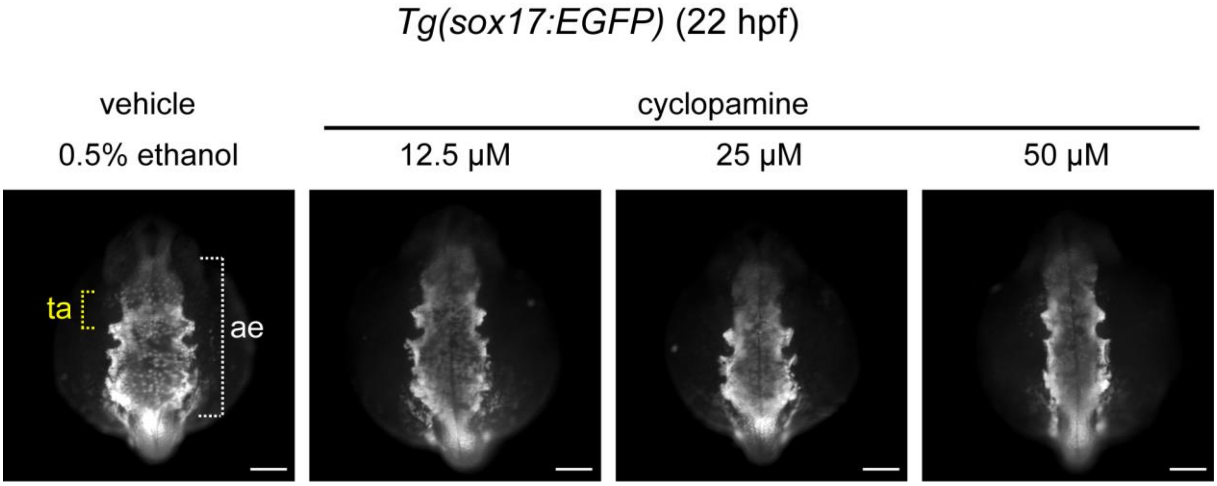


**Supplementary Figure 5. Inhibition of Shh signaling during somitogenesis affects anterior endoderm formation.** Transgenic *Tg(sox17:EGFP)* embryos were treated from 10 to 26 hpf with cyclopamine, an inhibitor of Shh signaling, and GFP reporter expression was analyzed by immunofluorescence staining at 22 hpf to assess formation of the anterior endoderm (ae). The prospective region of thyroid anlage (ta) formation within the anterior endoderm is highlighted by yellow brackets. Phenotypic comparison between treated and control embryos (0.5% ethanol) revealed concentration-dependent reduction of anterior endoderm in cyclopamine-treated embryos. Note the diminished mediolateral size of the anterior endoderm in cyclopamine-treated embryos. Dorsal views are shown, anterior is to the top. Scale bars: 100 µm.
